# DNA Sequence Preference for *De Novo* Centromere Formation on a *Caenorhabditis elegans* Artificial Chromosome

**DOI:** 10.1101/2020.04.12.037994

**Authors:** Zhongyang Lin, Karen Wing Yee Yuen

## Abstract

Centromeric DNA sequences vary in different species, but share common characteristics, like high AT-content, repetitiveness, and low, but not no, transcriptional activity. Yet, neocentromeres can be found on non-centromeric, ectopic sequences, suggesting that centromeres can be established and maintained epigenetically. In contrast, canonical centromeric DNA sequences are more competent in *de novo* centromere formation on artificial chromosomes (ACs). To determine if specific DNA sequence features are preferred for new centromere formation, we injected different DNA sequences into the gonad of a holocentric model organism, *Caenorhabditis elegans*, to form ACs in embryos, and monitored mitotic AC segregation. We demonstrated that AT-rich sequences, but not repetitive sequences, accelerated *de novo* centromere formation on ACs. We also injected fragmented *Saccharomyces cerevisiae* genomic DNA to construct a less repetitive, more complex AC that can propagate through generations. By whole-genome sequencing and *de novo* assembly of AC sequences, we deduced that this AC was formed through non-homologous end joining. By CENP-A^HCP-3^ chromatin immunoprecipitation followed by sequencing (ChIP-seq), we found that CENP-A^HCP-3^ domain width on both the AC and endogenous chromosomes is positively correlated with AT-content. Besides, CENP-A^HCP-3^ binds to unexpressed gene loci or non-genic regions on the AC, consistent with the organization of endogenous holocentromeres.

## INTRODUCTION

Genetic information is stored in DNA and packaged into chromosomes in all eukaryotic species. To ensure normal functioning of each individual cell and survival of the organism, the integrity of the genome must be maintained by duplicating chromosomes accurately, and separating them precisely into each daughter cell during each cell division. Errors in chromosome segregation can lead to aneuploidy and chromosomal instability.

Centromere function is essential for facilitating faithful chromosome segregation during mitosis, where the kinetochores build on the centromere to connect to the mitotic spindles. Ectopic centromere formation could cause merotelic attachments, chromosome breakages and rearrangements. Neocentromeres in cancer cells and patients with chromosomal rearrangements are usually found on non-centromeric sequences (1), suggesting that centromeres can be formed and maintained epigenetically (2). However, how and why neocentromere formation was initiated at ectopic regions is challenging to address, as neocentromeres can only be identified after they were formed (3).

On the other hand, constructing artificial chromosomes (ACs) in many species involves the use of canonical centromeric DNA sequences, like in budding yeast, fission yeast and human cells (4-7). This limits the options of sequences that can be tested when studying centromere formation. However, *de novo* centromeres in *C. elegans* embryos do not appear to have stringent sequence requirement (8), which allows us to test the efficiency of different DNA sequence features, such as variations in AT-content, repetitiveness and transcription levels, in *de novo* centromere formation.

When exogenous, naked DNA is injected into the *C. elegans* gonad, they could be concatemerized into high molecular weight (HMW) DNA arrays in oocytes, followed by chromatinization and *de novo* centromere establishment in the embryonic cells (8-10) (also refer to the back-to-back submitted manuscript). By co-injection of to-be-tested DNA sequences with a low amount of tandem LacO DNA repeats into a worm strain that expresses GFP::lacI in the germline and embryos, we can monitor the segregation process of newly formed ACs using live-cell time-lapse imaging (8). In this study, we demonstrated that high AT-content sequences, but not repetitive sequences, accelerated the process of *de novo* centromere formation on newly formed ACs.

The newly formed ACs can also be transmitted to subsequent generations, which allows us to expand the isogenic population to study the holocentromere positions and sequences biochemically on a propagated artificial chromosome. Here, we generated a stably propagated AC in *C. elegans* by co-injection of enzyme-digested budding yeast genomic DNA and several gene markers that are driven by different *C. elegans* promoters with different expression timings. We sequenced the whole genome of this *C. elegans* strain containing a complex AC, and *de novo* assembled the AC sequences. We mapped CENP-A^HCP-3^ binding sites on this AC, and demonstrated that *de novo* centromeres on the AC and endogenous centromeres share some common characteristics, such as high AT-content, and unexpressed gene loci or intergenic regions. Our study has shed light on the regulation of *de novo* holocentromere formation in a chromosome-wide landscape.

## MATERIAL AND METHODS

### METHODS

#### Plasmid construction

The 52% AT DNA fragment was PCR amplified from the genomic DNA of budding yeast *Saccharomyces cerevisiae* using the primers AT-52-F (5’-ATAATCTGCGACGAAGCTATGC-3’) and AT-52-R (5’-AAAGGAAGGCTCAATGACGAAT-3’). The 66% AT DNA fragment was amplified from the genomic DNA of budding yeast using the primers AT-66-F (5’-AAACGACAGCGAAGATAACG-3’) and AT-66-R (5’-TTGGGAGCCTATTGTGAAGT-3’). The 74% AT DNA fragment was amplified from the mitochondrial DNA of budding yeast using the primers AT-74-F(5’-GCAACTGAACTACCAGCAAG-3’’) and AT-74-R(5’-CCTGAATGTGCCTGAATAGA-3’). The amplified DNA fragments (∼5 kb) with different AT-contents were cloned into plasmid pMD-19T (2694 bp) by TA cloning to make pAT-52%, pAT-64% and pAT-74% (∼7 kb in total), respectively.

WYYp228 consists of a drug selection marker *Prsp-27::NeoR::unc-54 3’ UTR*, and two fluorescent marker genes, *Pmex5::GFP::tbb-2 3’ UTR* and *Pmyo-3::mCherry::unc-54 3’ UTR*. The promoter and 3’ UTR sequences were amplified from the genomic DNA of *C. elegans* strain EG8079, and cloned into the vector. The marker genes, NeoR-GFP-mCherry (NGM), are PCR amplified from WYYp228 using primers M13-47 (5’-CGCCA GGGTT TTCCC AGTCA CGAC-3’) and RV-M (5’-GAGCG GATAA CAATT TCACA CAGG-3’), and used for co-injection. Progenies from injected N2 worms were selected on G418-containing plates (1.5 mg/mL), and the individual survivors were singled out and further passed on for 10 generations.

#### Preparation of sheared or digested genomic DNA for co-injection

Salmon sperm DNA (Sigma-Aldrich) was sheared by Coveris M220 to a mean size of 5 kb. Yeast genomic DNA was isolated by glass beads method (11), followed by double digestion with restriction enzymes, *AfaI* and *PuvII*. All the fragmented genomic DNA were purified by Qiagen PCR purification kit.

#### Time-lapse live-cell imaging and AC segregation assay

To maintain a stable expression level of GFP::LacI, strain OD426 (Table 2) was maintained at 22°C. Artificial chromosomes (ACs) were visualized by injecting DNA containing LacO tandem repeats (8,12). Purified plasmid p64xLacO or its linear form (L64xLacO, digested by AfaI) (100 ng/μl) was injected into gonads of young adult *C. elegans* as reported by Yuen et al (8). To generate ACs with different AT-contents, pAT-52%, pAT-66% and pAT-74% were individually co-injected with p64xLacO in a ratio of 10:1. To increase the sequence complexity of the AC, sheared salmon sperm DNA, with a mean size of 5 kb, was co-injected with L64xLacO in a ratio of 10:1. The ratio of yeast genomic DNA (150 ng/ul) and NGM PCR product (0.5 ng.ul) is 300:1.

**Table 1.** Comparison of the assembly of the AC by directly assembling the whole genome (endogenous chromosomes and the AC), or by assembling reads that are excluded worm genomic sequences, as outlined in Figure 4 A.

|  | **Direct assembly of**<br>**AC from WGS reads** | **Assembly of AC**<br>**after filtering out**<br>**worm genomic**<br>**sequences** |
| --- | --- | --- |
| **Contig number** | 65 | 50 |
| **Total length** | 10,777,425 bp | 10,813,732 bp |
| **N50** | 418,241 bp | 691,347 bp |
| **Longest contig** | 1,487,696 bp | 1,706,458 bp |

**Table 2.** Worm strains used in this study.

| Strain name | Genotype | Reference |
| --- | --- | --- |
| N2 | Wild-type |  |
| OD426 | *ItIs37 [pie-1p::mCherry::his-58; unc-119(+)] IV.*<br>*mels1 [pie-1p::GFP::LacI]* | (19) |
| WYY35 | *Ex [Prps-27::NeoR::unc-54 3'UTR; Pmex5::GFP::tbb-2 3'UTR;*<br>*Pmyo-3::mCherry::unc-54 3'UTR; S. cerevisiae Afal, Pvull-*<br>*digested genomic DNA]* | This study |

Injected worms were recovered on OP50-seeded plates for 5-8 hours before time-lapse live-cell imaging. 3-4 worms were then dissected in 2 μl M9 buffer to release embryos. Embryos were mounted on a freshly prepared 2% agarose pad and the slide edges were sealed with Vaseline. Live images were taken with a Carl Zeiss LSM710 laser scanning confocal microscope with a 16 EC Plan-Neofluar 40x Oil objective lens and PMT detectors. Stacks with 17 x 1.8 μm planes were scanned for each embryo in a 3x zoom and a 1-minute or 30-second time interval, with 1.27 μs pixel dwell and 92 μm pinhole. Laser power for 488 nm and 543 nm was set at 5.5% and 6.5%, respectively.

To determine the AC segregation rates, every dividing cell that contains at least one AC was counted as one sample. Each division was categorized as either containing at least a segregating AC, or containing all non-segregating AC(s). Segregating ACs were defined as those that aligned with the metaphase plate and segregated equally with endogenous chromosomes during anaphase, or those that align to metaphase plate but do not segregate equally. Non-segregating ACs were defined as those that remained in the cytoplasm and did not separate in mitosis. The segregation rate was calculated as the number of dividing cells containing segregating ACs over the total number of dividing cells containing ACs. Among segregating ACs, those with bridges were referred to ACs that attempted to segregate, but the segregation process was incomplete and ACs were lagging during anaphase.

#### Fluorescence in situ hybridization (FISH)

1 μg of *S. cerevisiae* (SY15, a single circular chromosome strain derived from BY4742 (13)) genomic DNA were labeled by nick translation to generate green fluorescent probes using Atto488 Nick Translation Labeling Kit (Jenabioscience) and purified by silica-gel membrane adsorption (Qiagen PCR Purification Kit), according to the kit protocols. Adult worms were dissected in 2 μl of 1X egg buffer on a coverslip (25 mM HEPES, pH 7.3, 118 mM NaCl_2_, 48 mM KCl, 2 mM CaCl_2_, 2 mM MgCl_2_) to release embryos. Coverslips were placed in liquid nitrogen followed by freeze-cracking and then fixed in -20°C methanol for 30 minutes. Slides were washed twice in 1X PBS and fixed in PBS with 4% formaldehyde for 5 minutes. Slides were washed twice in 1XPBS and permeabilized by PBST (0.5% Triton X-100) for 15 minutes. Slides were washed with 2X SSC for 5 minutes, followed by incubation with 2X SSC-RNase (2 mg/mL) at 37°C for 45 minutes. Samples were resuspended in the Hybridization mixture (10% dextran sulfate, 2X SSC, 50% formamide, 160 ng/µl sheared salmon sperm DNA, 8 ng/ul probe DNA) at 90°C for 5 minutes to denature the samples, and followed by incubation in humidified chamber at 37°C overnight. On the next day, samples were washed 3 times with prewarmed 2X SSC/50% formamide for 5 minutes at 42°C, followed by three 5-minute washes with 2X SSC at 42°C and a 10-minute wash in 1X SSC at 42°C. Wash buffer was removed and samples were stained by DAPI (1μg/mL) at room temperature for 10 minutes, followed by a 10-minute wash with PBST. Mounting was performed using ProLong gold antifade reagent (Life Technologies). The slides were sealed with nail polish and stored at -20°C before imaging.

#### Immunofluorescence (IF) staining

Embryos were freeze-cracked after dissection of adult worms and fixed in -20°C methanol for 30 minutes. Embryos were then rehydrated in PBS (137 mM NaCl, 2.7 mM KCl, 4.3 mM Na_2_HPO_4_, 1.4 mM KH_2_PO_4_) for 5 minutes and blocked by AbDil (4% BSA, 0.1% Triton-X 100 in PBS) at room temperature for 20 minutes. Primary antibody incubation, using rabbit (Rb)-anti-AIR-2 (1:500, a gift from Arshad Desai Lab), Mouse (Ms)-anti-LacI (1:250, Millipore 05-503), was performed at 4°C overnight. Slides were washed with PBST 3X 10 minutes. The slides were then incubated with goat-anti-Ms-IgG FITC secondary antibody (1:100,000; Jackson ImmunoResearch Laboratories, 115-096-062) and goat-anti-Rb-IgG Alexa 647-conjugated secondary antibody (1:100,000; Jackson ImmunoResearch Laboratories, 111-606-045) at room temperature for 1 hour, followed by DAPI (1μg/mL) staining for 15 minutes. Mounting was done using ProLong gold antifade reagent (Life Technologies).

Images were acquired from Zeiss LSM 780 upright confocal microscope with a Plan-Apochromat 40x/1.4 Oil DIC M27 objective and PMT detectors. Embryos were captured as z stacks with 30-35 z-sections and a z-step size at 0.4 μm and 3.15 μs of pixel dwell time. Stacks were scanned for each embryo in a 4x zoom. DAPI, FITC, and Alex647 channels were scanned with 32 μm pinhole and the images were saved in 16 bits format.

#### MinION library preparation

The genomic DNA from worm strain WYY35 (Table 2) at L1 stage was sequenced using Oxford Nanopore sequencing technology. Briefly, high molecular weight DNA was sheared with a g-TUBE (Covaris) to an average fragment length of around 8 kb. The sheared DNA was repaired using the FFPE Repair Mix, according to the manufacturer’s instructions (New England Biolabs). 0.4X Ampure XP beads (Beckman Coulter) was used to exclude short DNA fragments. The DNA ends were blunted and an A overhang was added with the NEBNext End Prep Module (New England Biolabs). Prior to ligation, the DNA solution was cleaned up again by 1X Ampure XP beads. The adapter was ligated to the end-repaired DNA using Blunt/TA Ligase Master Mix (New England Biolabs). The final library was eluted from 0.4X Ampure XP beads after washing 2 times by Adapter Bead Binding buffer (SQK-LSK108 Ligation Sequencing Kit 1D). Two R9.4 Flow-cells were used to sequence the DNA. The MinKNOW software (version 1.5.12) was used to control the sequencing process, and the raw read files were uploaded to the cloud-based Metrichor EPI2ME platform for base calling. Base-called reads were downloaded for further processing and assembly.

#### Mi-Seq library preparation

Genomic DNA from worm strain WYY35 was sheared by Covaris M220 using the default setting for fragment length of 500 bp. About 1 μg of genomic DNA from WYY35 were used for library construction using NEBNext® Ultra™ II DNA Library Prep Kit. The constructed library was run on 1% agarose gel electrophorese for size estimation, and quantified using NEBNext® Library Quant Kit for Illumina®. The library DNA was then diluted to 4 nM and denatured by 0.2 N NaOH for 5 minutes at 95°C, as described in MiSeq System Denatureand Dilute Libraries Guide. Lastly, 12 pM of denatured library DNA was loaded onto the reagent cartridge for the sequencing run. MiSeq sequencing was performed with the assistance from Dr. Zhao’s lab at Hong Kong Baptist University. Adapter trimming of the raw reads was performed using Trimmomatic (14).

#### Hi-seq library preparation (for ChIP-seq)

Library preparation and Illumina sequencing (Pair-End sequencing of 101 bp) were performed at the University of Hong Kong, Centre for Genomic Sciences (HKU, CGS). The 4 libraries (two replicates of Input and ChIP-ed DNA) were prepared based on the protocol of KAPA Hyper Prep Kit (KR0961 – v5.16). For each library, 0.8 ng of DNA was performed with reactions of end-repair, 3’ end A-tailing, and indexed adaptor ligation, followed by 16 cycles of PCR amplification reaction for library enrichment. After AMPure beads purification, each library was validated by Agilent Bioanalyzer, Qubit and qPCR for quality control analysis. The library was denatured and diluted to optimal concentration and applied in the cluster generation steps. HiSeq PE Cluster Kit v4 with cBot was used for cluster generation on the flow-cell. Illumina HiSeq SBS Kit v4 was used for Pair-End 101bp sequencing that runs on HiSeq 1500.

#### *De novo* genome assembly and evaluation

Adapter trimming of the raw reads was produced from Mi-Seq using Trimmomatic (14). All trimmed pair-end reads were used for *de novo* assembly by SPAdes in its default settings (Command: python2.7 ∼/SPAdes-3.5.0-Linux/bin/spades.py -t 24 -m 90 -k 21,33,55,77,99,127 –careful -1 AC_Trim_1P.fq -2 AC_Trim_2P.fq -o spades_Miseq_assembly) (15).

All MinION reads from 2 flow-cells, despite their quality, were combined and used for Canu nanopore assembly in its default settings (Command: canu -p gm_mini_merged_umpromo -d gm_mini_merged_umpromo genomeSize=20m useGrid=false overlapper=mhap utgReAlign=true correctedErrorRate=0.10 -nanopore-raw gm_mini_merged_umpromo-fiter.fa) (16). To compare with the assembly result using all MionION reads, high quality reads from the “Pass” category for MinION reads that have a quality score above “q6” were filtered based on a quality metric by Metrichor and used for assembly separately. High accurate paired reads generated from Mi-seq platform were re-mapped to the assembled contigs.

Small indels and misassembles were corrected by Pilon (17). The correcting process was repeated 3 times until the alignment identity to the reference genome no longer improved. Assemblies from MinION reads were evaluated by dnadiff in MUMmer3 (18) to align draft assemblies against the *C. elegans* reference genome (WS254). After each round of Pilon correction, the increase in sequence identity and aligned bases to the reference were compared.

#### ChIP-seq analysis

Paired-end reads from ChIP and input samples were mapped to the reference genome (WS245) using BWA-mem (19). Mapped reads from ChIP and input were used to call peaks and obtain read coverage per base using MACS version 1.4.2 under broad domain setting with P-value = 1e-3 cutoff. (Command: macs2 callpeak -t Chip_AC.bam -c InPut_AC.bam -f BAM -g 11072328 -n domain_ACv5 -B -p 1e-3 --broad --broad-cutoff 0.1 --fix-bimodal --extsize 100 --outdir ∼/jobs/chipseq_mapping/AC_masc_domains/domains). Log2 enrichment ratios (ChIP/Input) of each replicate were calculated using Deeptools software (20). To compare with published ChIP-chip data, previously reported CENP-A^HCP-3^ microarray data (21) were obtained from modENCODE. For comparing the correlation between ChIP-seq and ChIP-chip, and among ChIP-seq replicates, the average scores of each 1-kb bin over the entire genomic region were calculated. The Pearson correlation coefficients were calculated among all data sets and shown in scatter plots. The Spearman correlation coefficient were calculated among all data sets for plotting a heat-map graph. Hierarchical clustering was applied to the correlation matrix. CENP-A^HCP-3^ domains that are defined by MACS was plotted according to their width and AT% by ggplot2 in R. CENP-A^HCP-3^ motif was found by MEME Version 5.1.0 (Command: meme ./seqs-centered -oc meme_out -mod zoops -nmotifs 3 -minw 6 -maxw 30 -bfile ./background -dna - searchsize 100000 -time 5082 -revcomp -nostatus). Motif-motif similarity was measured by Tomtom (22) (Command: tomtom -no-ssc -oc. -verbosity 1 -min-overlap 5 -mi 1 -dist pearson -evalue -thresh 10.0 query_motifs db/WORM/uniprobe_worm.meme).

## RESULTS

### High AT-content, but not repetitive sequence, accelerates the ability of AC to acquire segregation competency

To determine the effects of AT-content on new centromere formation, a ∼5-kb insert with different AT-contents were PCR-amplified from *S. cerevisiae* genomic or mitochondrial DNA, with low (52%), medium (66%) or high (74%) AT-contents, and cloned into the pMD19-T vector backbone (2694 bp). The final constructs were called pAT-52%, pAT-66%, and pAT-74%, respectively. Individual plasmids were co-injected with the p64xLacO plasmid in a 10:1 ratio into the syncytial gonad of a *C. elegans* strain expressing GFP::LacI and mCherry::H2B. After 6 hours of microinjection, embryos were dissected from injected worms for live-cell time-lapse imaging. In 1-4-, 5-8- and 9-16-cell stage embryos, the segregation rates of ACs generated from pAT-52% and pAT-66% show no significant difference. However, the segregation rates of ACs made with pAT-74% are significantly higher than those of pAT-52% and pAT-66% (Figure 1 B). This finding indicates that ACs with this high AT-content sequence acquire segregation competency faster than those with low and medium AT-content sequences in 1-16-cell stage. At 17-32-cell stages, the segregation rates of ACs generated from the three AT-contents become similar. Our findings indicate that higher AT-content may accelerate the process of *de novo* centromere formation.

**Figure 1.**
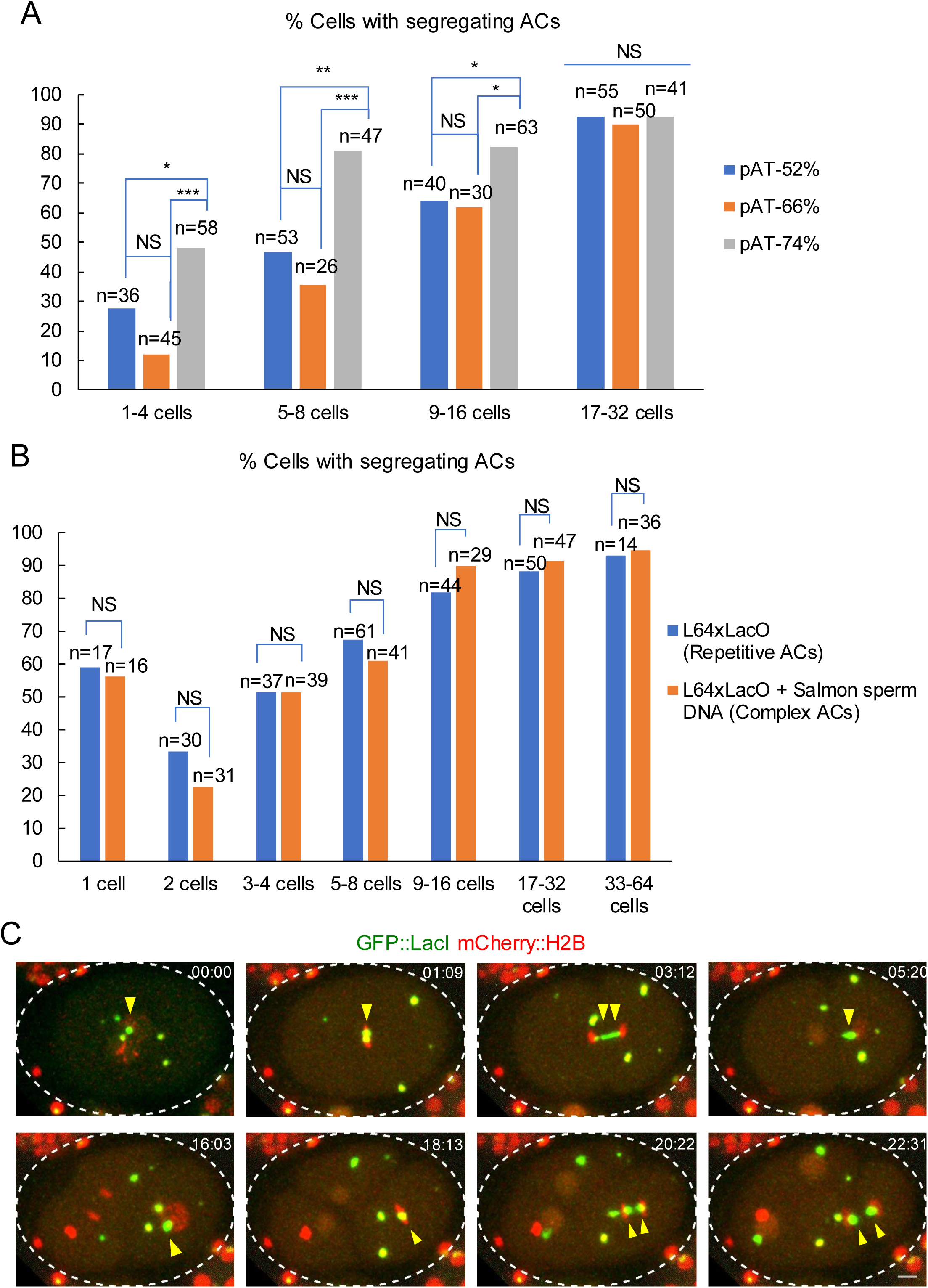
AC segregation and the relationship with injected DNA sequence characteristics. (A) Quantification of the cells with segregating ACs after injection of plasmid DNA with different AT-contents. pAT-52%, pAT-66% or pAT-74% was co-injected with the p64xLacO plasmid in a ratio of 10:1 to generate ACs with different AT-contents. All types of DNA for microinjection were adjusted to 100 ng/μL in total. The percentage of cells with segregating ACs among all dividing cells containing ACs was scored. The number of cells (n) analyzed was indicated. Blue arcs show comparisons among pAT-52%, pAT-66% and pAT-74% at each cell stage. Chi-square test was used to test for significance. *p < 0.05, **p < 0.01 and ***p < 0.001. NS means not significant. Although the segregation competency of all types of ACs improves over time, the difference of their segregation competency is more significant in early stage than in late-stage embryos. (B) Quantification of the percentage of cells with segregating ACs after injection of DNA with different sequence complexity. ACs with complex sequence context were generated by injection of a mixture of linear plasmid (L64xLacO) and sheared salmon sperm DNA (SS-DNA) (∼5 kb) in a concentration ratio of 1:10. The segregation rate of complex ACs were compared with that of repetitive ACs, which are generated by injecting just the linear plasmid (L64xLacO) (Refer to the back-to-back submitted manuscript). All types of DNA for microinjection were adjusted to 100 ng/μL. The percentage of cells with segregating ACs among all dividing cells containing ACs was scored. The number of cells (n) analyzed was indicated. Chi-square test was used to test for significance. *p < 0.05, **p < 0.01 and ****p < 0.0001. NS means not significant. (C) 5 hours after injection of L64xLacO, a representative embryo expressing GFP::lacI (green) and mCherry::H2B (red) and carrying multiple ACs is shown by live-cell time-lapse imaging. Yellow arrows points to the AC undergoing chromosome segregation from 1-cell stage to 4-cell stage. The time (mm:ss) is indicated on left top of images. These ACs mature from passive inheritance to autonomous segregation rapidly within a few cell cycles. Scale bar represents 5 µm.

To reveal if repetitive sequences benefit *de novo* centromere formation, we injected linearized p64xLacO sequence (L64xLacO), with or without sheared salmon sperm DNA. A representative embryo, which carries an AC, that is undergoing chromosome segregation from 1-cell stage to 4-cell stage is shown (Figure 1C). The addition of salmon sperm DNA (10:1 L64xLacO) allows the formation of ACs with a complex sequence context (23). However, AC segregation analysis shows no significant difference between repetitive and complex ACs, which suggests that repetitive sequences do not facilitate *de novo* holocentromere formation on ACs (Figure 1 C).

### Construction of a self-propagated artificial chromosome in *C. elegans* using sheared yeast genomic DNA

To understand how the foreign DNA microinjected to the *C. elegans* gonad is assembled into an artificial chromosome, we constructed an AC with more sequence complexity, and with sequences that can be distinguished from endogenous *C. elegans* sequences. Thereby, genomic DNA from another species was used. We sheared the budding yeast *Saccharomyces cerevisiae* (BY4742, a S288C-derivative strain) genomic DNA by restriction enzymes, AfaI (GT|AC) and PvuII (CAG|CTG), to generate blunt-ended short DNA fragments (50-8448 bp) (Figure 2 A). The agarose gel electrophoresis image of restriction enzyme-digested budding yeast genomic DNA confirms that the DNA length distribution is between ∼100-2000 bp (Figure 2 B).

**Figure 2.**
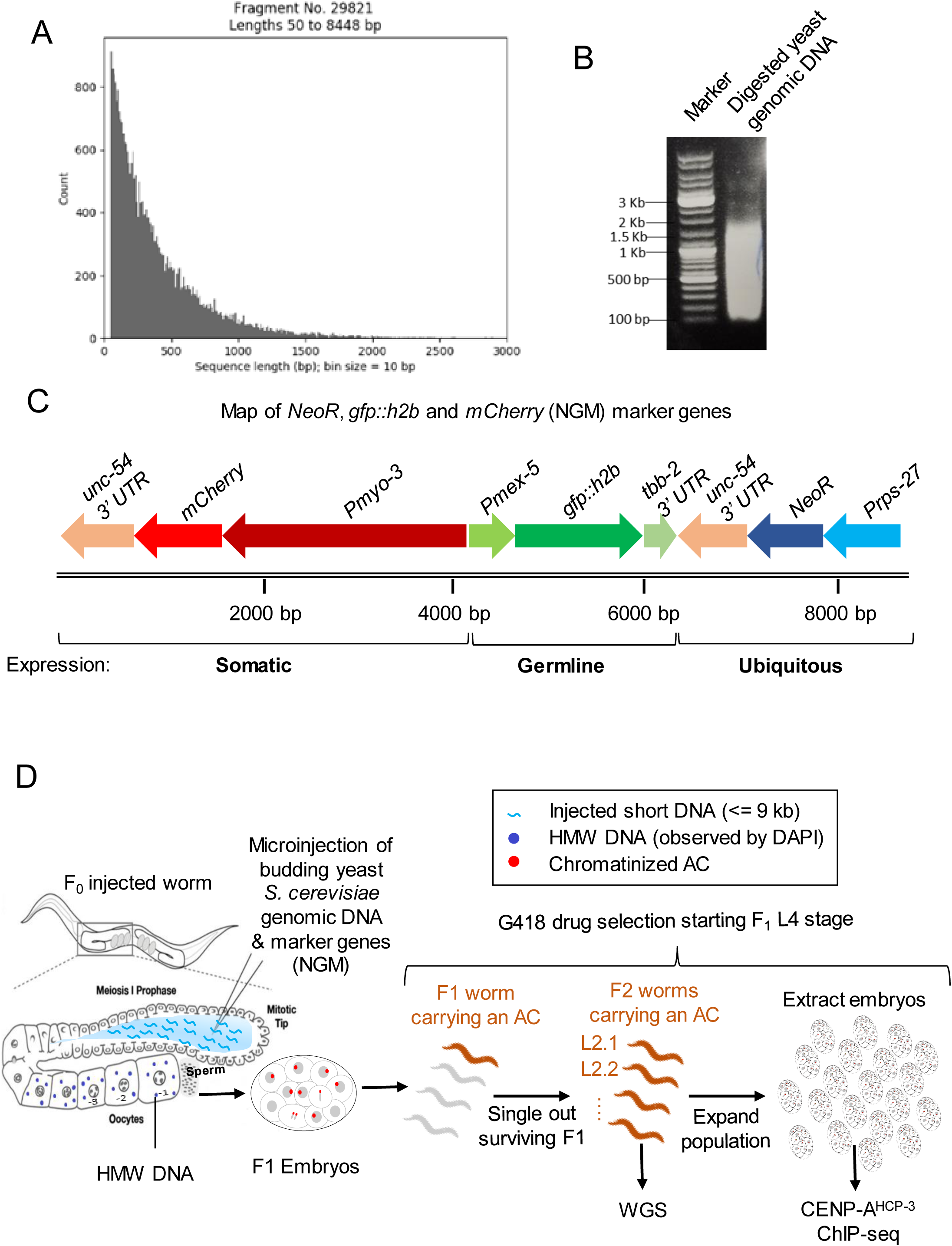
Construction of a complex, propagated artificial chromosome in *C. elegans* using fragmented yeast genomic DNA and marker genes. (A) Budding yeast *Sacchromyces cerevisiae* genomic DNA is digested with restriction enzymes AfaI (GT|AC) and PvuII (CAG|CTG). A histogram of *in silico* digestion is shown. (B) Genomic DNA was isolated from an S288C background strain, BY4742. An agarose gel image showing the sizes of enzyme-digested DNA (AfaI and PvuII). 150 ng/µl of this digested DNA is used for microinjection. (C) A schematic map of the co-injection markers, *<u>N</u>eoR, <u>g</u>fp::h2b*, and *<u>m</u>Cherry* (NGM), PCR-amplified from plasmid WYYp228. The ubiquitous antibiotics resistance gene (NeoR) is driven by *rps-27* promoter. *gfp::h2b* is driven by germline *mex-5* promoter, and mCherry is driven by somatic, body wall muscle *myo-3* promoter. The scale bar (in bp) is shown. A PCR product from the WYYp228 containing NGM (0.5 ng/µl) is used in the co-injection mix. (D) Schematic of the experimental approach used to generate a complex, propagated artificial chromosome in *C. elegans*.

To harvest enough worms and embryos carrying an artificial chromosome for DNA sequencing and subsequent chromatin immunoprecipitation analysis, respectively, we applied antibiotics selection method for enrichment (24). A neomycin resistance gene (NeoR) under the control of the *C. elegans* ubiquitous *rps-27* promoter and *unc-54 3’ UTR*, a germline expression marker *Pmex-5::GFP::H2B::tbb-2 3’ UTR*, and a somatic expression marker *Pmyo-3::mCherry::unc-54 3’ UTR* are cloned onto a vector, and we called these three gene markers <u>N</u>eoR-<u>G</u>FP-<u>m</u>Cherry, or NGM for short (Figure 2 B). We mixed restriction enzyme-digested yeast genomic DNA (150 ng/μl), with a very low amount of the NGM marker (0.5 ng/μl), for co-injection into wild-type (N2) *C. elegans* worms.

F1 progenies from injected N2 worms were selected on G418-containing plates, and individual survivors were singled out and further passed on for 10 generations. In *S. cerevisiae*, there is a single cluster of rDNA, comprising of approximately 150 copies, located on chromosome XII. This cluster covers about 60% of chromosome XII and about 10% of the whole genome. The copy number of NGM markers and yeast rDNA from two sublines (L2.1 and L2.2), which originated from the same F1 progeny, with stable body wall expression of mCherry, was quantified by real-time PCR. The Ct value for mCherry and rDNA PCR were then normalized to that of a unique worm gene locus on chromosomes II (9565795:9565963). This unique worm genomic locus has 2 copies in a diploid genome in a *C. elegans* cell. The ratios of mCherry and rDNA to this worm locus were used to deduce the copy number of mCherry and rDNA per cell. Subline L2.1 is estimated to contain ∼42 copies of mCherry and ∼302 copies of yeast rDNA which was close to two times of the copy number of rDNA per haploid yeast genome (25). We chose subline L2.1 (WYY35) for further analysis, as L2.1 contains a lower mCherry copy number than subline L2.2, suggesting that L2.1 AC sequences are more complex than that in L2.2.

To observe the AC, we performed DAPI staining, immunofluorescence analysis and fluorescent *in situ* hybridization (FISH) in oocytes and embryos in subline L2.1 after multiple generations of propagation. DAPI staining shows that the AC was maintained as an independent chromosome that is well separated from the 6 compact endogenous chromosomes in diakinesis oocytes (Figure 3B top and middle). Immunofluorescence analysis shows that AIR-2 is absent on the propagated AC, suggesting that it is a monosomic, univalent chromosome (Figure 3B top). FISH analysis using probes generated from yeast genomic DNA further confirms the AC as an extra-chromosomal array in diakinesis oocytes (Figure 3 B middle).

**Figure 3.**
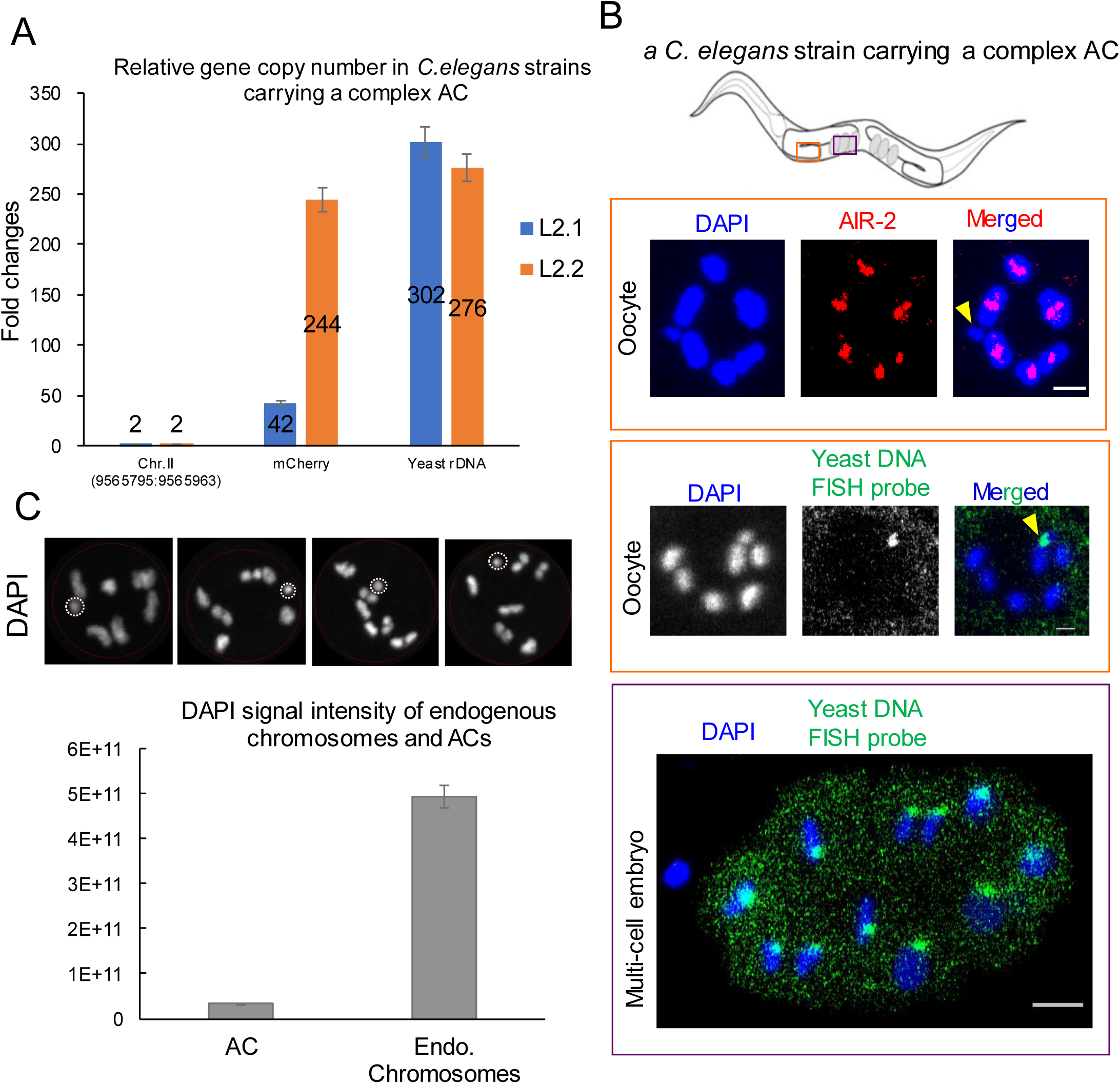
Isolating *C. elegans* strains carrying an AC with a low copy number of marker genes, and an estimation of the AC size by cytology. (A) Quantification of an endogenous gene on Chromosome II (9565795:9565963), mCherry marker, and yeast rDNA copy number in strains with an AC by real-time PCR. L2.1 and L2.2 are different F2 progenies from the same F1 produced by an injected worm. Error bars indicate 95% confidence interval (CI) for the mean. (B) The propagated AC (arrowhead) was stained uniquely by the FISH probes made from yeast genomic DNA, and was lack of AIR-2 signal in the oocytes. In the multi-cell embryo, the propagated ACs aligned at the metaphase plate and segregated with endogenous chromosomes in anaphase. (C) Estimation of AC DNA size by DAPI staining. The DAPI signal intensity on the 6 endogenous chromosomes corresponds to 400 Mb (replicated diploid with 100-Mb genome size). The DAPI signal intensity on the propagated AC, assuming to be replicated and in monosomy, is used to calculate the size of this AC. ACs and endogenous chromosomes from 4 individual oocytes were used for quantification. The bar chart shows the DAPI signal intensity on the AC and on all the endogenous chromosomes. The DAPI signal on all endogenous chromosomes is about 15 folds of that on AC. Error bars indicate 95% confidence interval (CI) for the mean. The AC size (1n) was estimated to be around 13 Mb.

Besides, this yeast genomic DNA FISH signal on this AC in multi-cell embryo shows that the AC aligns at metaphase plates and segregates with endogenous chromosomes in anaphase (Figure 3B bottom), suggesting that this AC segregates normally during mitosis. However, we found that under a non-selective condition, the frequency of inheriting the AC to the next generation is about 60%, based on the presence of somatic body wall mCherry fluorescence. This result suggests that while the AC is quite mitotically stable, the unpaired AC tends to be lost in meiosis, similar to the loss rate of the X chromosome in the *him-8 null* background where two homologous X chromosomes did not paired in meiosis (26).

The DAPI signal of the AC was compared to that from the 6 bivalent endogenous chromosomes in diakinesis oocytes (Figure 3C), which contain replicated (2 copies) diploid genome (a total of ∼400 Megabases). Based on the assumption that the propagated AC is fully replicated in oocytes and that the condensation of the AC and endogenous chromosomes are comparable, the size of the AC was estimated to be about 13 Mb (Figure 3C), as calculated according to the equation: AC size (Mb) = DAPI intensity of AC X 400 Mb / (DAPI intensity of Endogenous Chromosomes X 2 sister chromatids).

### DNA sequencing of a *C. elegans* strain with a complex artificial chromosome

Genomic DNA from strain L2.1 (WYY35) was sequenced directly by nanopore MinION and Mi-seq without PCR to avoid PCR bias during library preparation. Raw data produced from two MinION flow-cells were base-called by Metrichor. Two R9.4 flow-cells ran for 48 hours to produce 1,442,172 base-called reads in total, with an average read length of 9,804 bases and N50 of 8,168 bases (Figure S1; Table S1). The average yield per flow-cell was about 5.0 Gigabases (Table S1). Plotting read length against base-call quality showed no correlation, indicating that there was no quality bias of the read length (Figure S1.C and D). All reads produced from 2 flow-cells were combined, and the read quality was evaluated by aligning to the *C. elegans* reference genome (WS245). Alignment percentage identity was highly correlated to the read quality (Pearson, r = 0.75), while no length bias was observed (Pearson r = 0.04) (Figure S2.A and B). Pass reads, filtered by Metrichor (Phred quality above 6), showed higher mappability and a lower error rate when compared with All reads. 79.19% of All reads and 88.49% of Pass reads were aligned to the reference genome by Graphmap (27), with a general error rate of 20.73% and 18.87%, respectively (Table S2).

### *De novo* genome assembly of the *C. elegans* strain with a complex artificial chromosome using MinION and Illumina sequence reads

All reads and Pass reads from MinION were separately assembled by Canu pipelines (16). Assembly from All reads had slightly lower contigs number and slightly higher N50 than assembly from Pass reads. The alignment of the assembled contigs covered 99.86% of the reference worm genome, without any large gap (Table S3; Figure S3). 241 contigs were generated from All reads with N50 of 1,487,696 bp. 145 of the contigs containing 103,389,851 bases (86.77% of the assembled contigs) with significant homology to 99.86% of the *C. elegans* reference genome, suggesting that the remaining 13.33% of the assembled contigs, corresponding to ∼15 Mb, do not belong to the reference worm genome (Table S2). 65 contigs, containing 10,777,425 bp, were homologous to budding yeast genomic sequence injected to generate the complex artificial chromosome.

The continuity of *de novo* assembly result of MinION reads by Canu was significantly higher than the assembly of Mi-seq reads using SPAdes, which had a N50 value of only 22,768 bp. However, assembly from the Mi-seq reads had higher 1-to-1 identity to the reference worm genome, 99.94%, as compared to 95.84% of the Canu assembly, indicating that Mi-seq reads had higher sequence accuracy (Table S3). Thus, Mi-seq reads were only used for polishing the Canu assembly. The percentage identity of Canu contigs to the reference worm genome was improved to 99.79% after polishing using the Mi-seq sequence data by applying Pilon (17) 3 times (Table S3).

### *In silico* filtering improves the *de novo* assembly result of the AC

From direct Canu assembly of All reads, we found that one contig (tig93) from worm chromosome II, also partially aligned to some yeast sequence. This hybrid contig was assembled through the common sequence of *myo-3* promoter in both *C. elegans* genome and NGM markers (Figure S4A). MinION reads re-mapped to this contig shows a low coverage around the *myo-3* promoter, indicating that this region was misassembled (Figure S4B). This is because the 2.5 kb *myo-3* promoter is longer than the parameter used in Canu assembly program in searching for a minimum 500-bp overlapping among reads (16,28).

Therefore, we used *in silico* filters to isolate reads of the artificial chromosome from the whole genome. Bacterial contaminated reads were first removed by aligning to the bacterial genome found in a pilot assembly (Figure 4, “Bacteria Filter”). The drug-resistant bacteria, *Stenotrophomonas maltophilia*, could attribute to the bacterial contamination on the G418 plates used to culture the nematodes, because the two *S. maltophilia* contigs are free from yeast or worm sequence. Then reads that aligned to the *in silico* digested yeast genome by two different algorithms, Graphmap and minimap2, were isolated (Figure 4, “NGM & Yeast Filter”) and combined with reads that did not align to the *C. elegans* genome (Figure 4, “Worm Genome Filter”). In “Worm Genome Filter”, reads were filtered through the *C. elegans* whole genome sequence, and one more time through the flanking regions (∼ 10kb) of worm promoters and 3’ UTRs in NGM marker, to improve the removal efficiency. All these isolated reads were merged, followed by removing the duplicates. After the Canu assembly pipeline, contigs were further aligned to the *C. elegans* reference genome and filtered out if they have > 95% identity (Figure 4, “Remove Worm Contigs”). The total length of the assembled AC after *in silico* filtering was extended from 10,777,425 bp to 10,813,432 bp. By assembling reads after *in silico* filtering, contig N50 of the AC also improved from 418,241 bp to 691,347 bp, together with a reduction in contig number (from 65 to 50) (Table 1).

**Figure 4.**
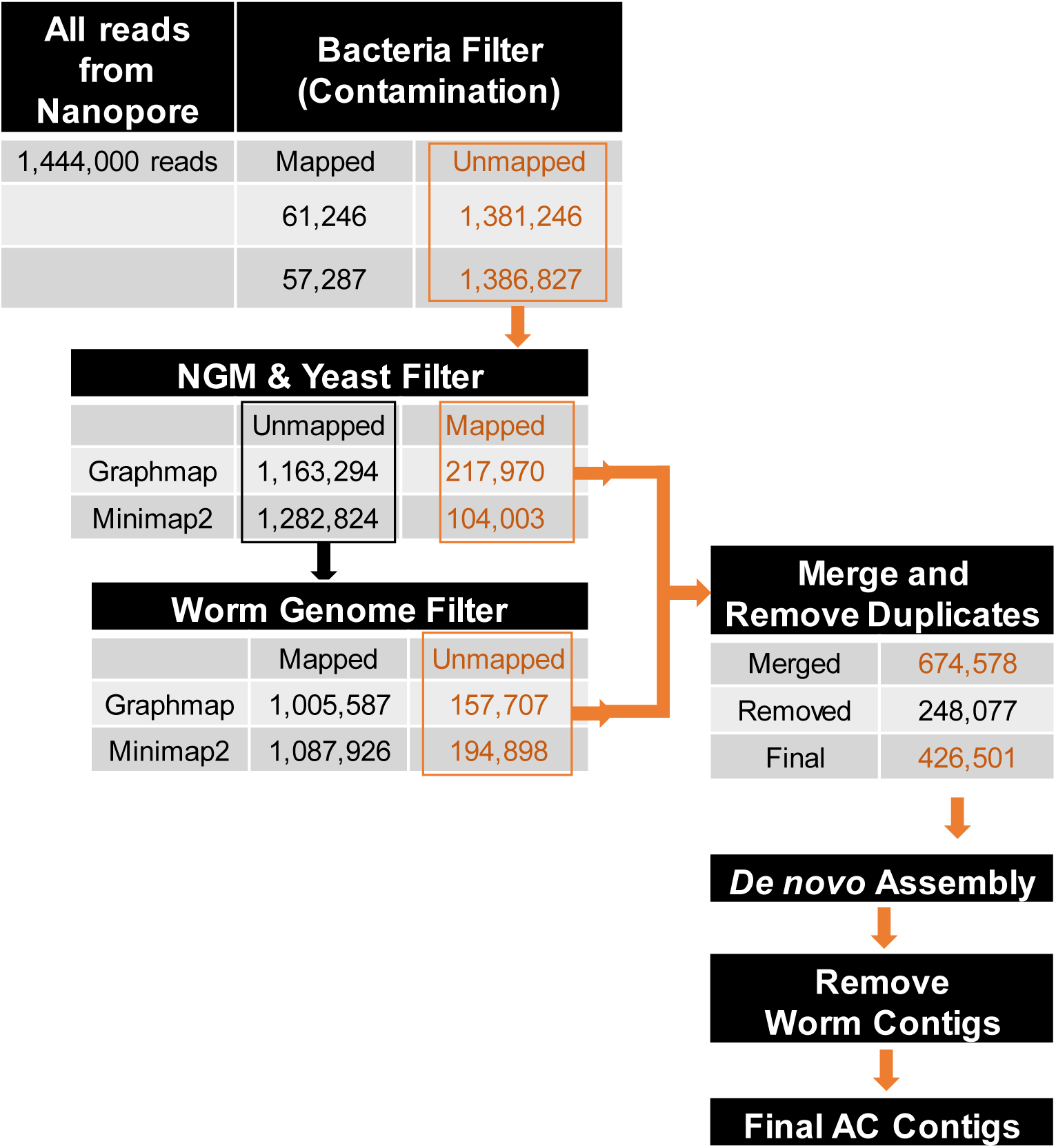
Workflow for isolating AC reads from the whole *C. elegans* genome sequencing (WGS). Contaminated bacterial sequence were first filtered out (contamination filter). All reads were then aligned to the *in silico* digested yeast genome sequences (NGM-yeast filter). The mapped reads from “NGM-yeast filter” were combined with unmapped reads from “Worm genome filter”. Two different algorithms, Graphmap and minimap2, were used for the mapping in parallel. Then, the isolated reads were merged, followed by removing duplicates. *De novo* assembly by Canu were then carried out. Low quality reads that belong to the *C. elegans* genome, but escaped from the *in silico* filters also went through the Canu assembly pipeline. The assembled worm contigs were filtered out after contigs assembly by blastn (Figure 4, Remove worm contigs).

### *De novo* assembly result suggests that this AC is formed by non-homologous end joining

A comparison between the AC sequence and the yeast genome using dnadiff showed that while this AC size is comparable to that of *S. cerevisiae* genome size, this AC consists of ∼30.9% yeast genomic sequence (which are used multiple times). Mummer alignment of the assembled AC contigs to the yeast reference genome (S288C) showed that the sequence of each contig was a random combination of short DNA fragments from each yeast chromosome (Figure 5A). The NGM co-injection markers were found to be interspaced as single copies in between yeast genomic DNA, but not concatemerized. For example, on the largest contig (tig8258), NGM markers and some of the incomplete NGM fragments were interspaced in between the yeast genomic DNA (Figure 5C). From the self-alignment of tig8258, we found that the same fragmented yeast sequences appeared multiple times on the assembled AC, but in different organizations. However, we have not identified obvious tandem repeats (Figure S5A).

**Figure 5.**
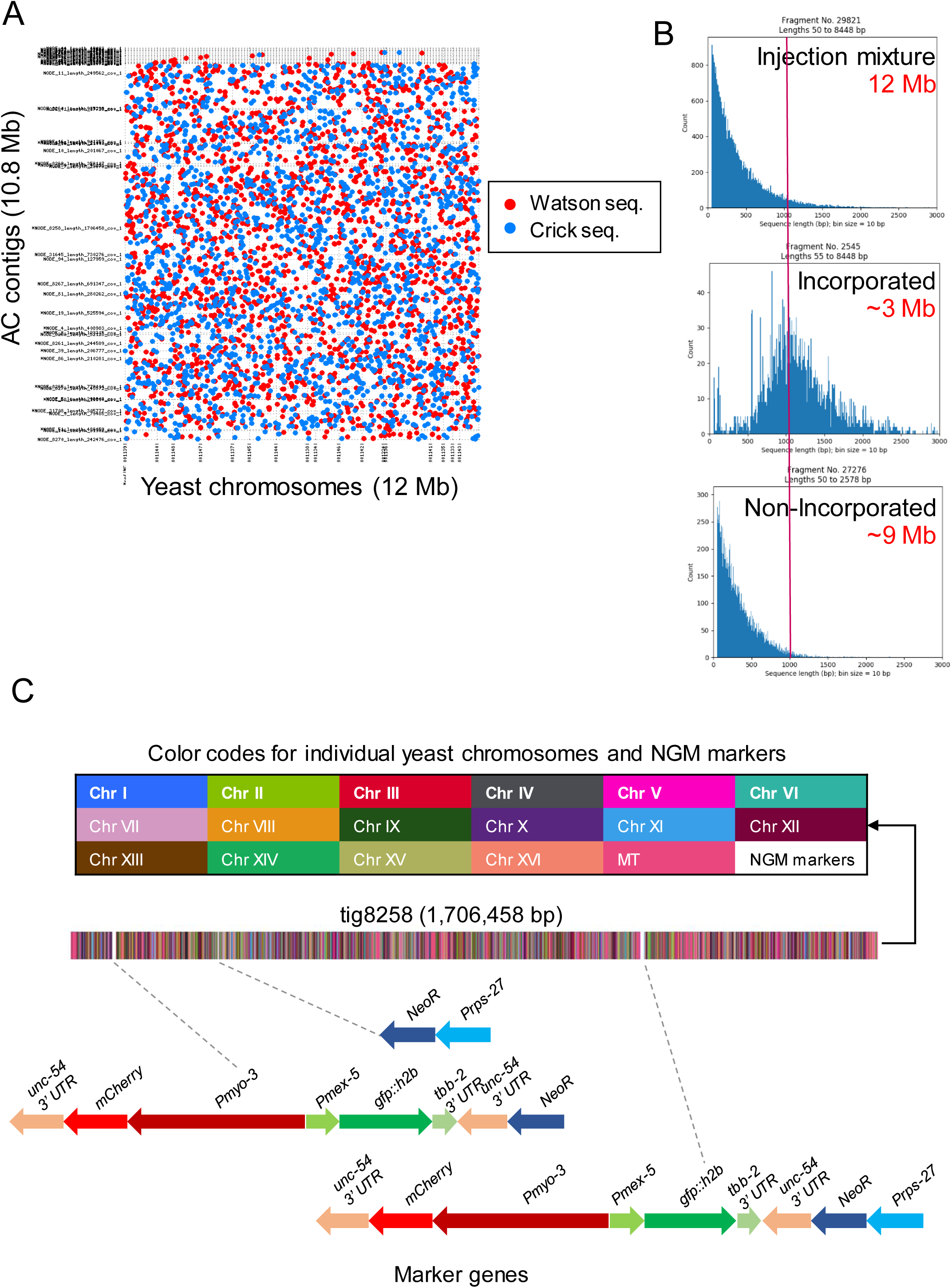
*De novo* assembly result suggested that this AC is formed by random non-homologous end-joining of the injected fragmented yeast genomic DNA and the PCR product from the marker plasmid. (A) Mummer alignment of assembled AC contigs to the yeast genome (Chromoosme I to XVI) shows that the sequence of each contig is a random combination of short DNA fragments from different yeast chromosomes. Red dots: 5’ to 3’ direction (Watson strand in SGD https://www.yeastgenome.org/seqTools#gene); blue dots: 3’ to 5’ direction (Crick strand in SGD). (B) Histogram plots of the DNA fragment lengths from *in silico* AfaI- and PvuII-digested budding yeast genome (as in Figure 2 B right), sequences incorporated into the AC (corresponding to larger fragments, comprising of ∼3 Mb) and sequences that did not incorporate into the AC (corresponding to smaller fragments, comprising of ∼9 Mb). The red line indicates the fragment length of 1 kb. (C) A schematic of distribution of yeast sequence fragments and the co-injection markers NGM (*Prps-27::NeoR::unc-54 3’ UTR; Pmex5::GFP::tbb-2 3’ UTR; Pmyo-3::mCherry:: unc-54 3’ UTR*) on the largest assembled contig tig8258. DNA fragments belong to individual yeast chromosomes and NGM marker genes are indicated by different colors.

### Identification of CENP-A^HCP-3^ loading patterns on a propagated AC

As cross-linked ChIP-microarray (ChIP-chip) against CENP-A^HCP-3^ on *C. elegans* endogenous chromosomes has been reported previously (21), we evaluated the reproducibility among our biological replicates of CENP-A^HCP-3^ ChIP-seq, and between previous reported CENP-A^HCP-3^ ChIP-chip results (1 kb bins). The Pearson correlation and Spearman correlation between ChIP-seq replicates were generally high (median r = 0.83 and 0.84, respectively), indicating that our ChIP-seq experiments against CENP-A^HCP-3^ were reproducible (Figure S6A and S6B). The cross-platform correlation between pairs of replicates of ChIP-chip and ChIP-seq were also high, with median r > 0.81, indicating that CENP-A^HCP-3^ ChIP-seq results corroborate previous CENP-A^HCP-3^ ChIP-chip results (Figure S6A and S6B). A representative genome browser view on 0.5 Mb on chromosome I for CENP-A^HCP-3^ ChIP-seq and ChIP-chip (21) showed similar patterns and domains of CENP-A^HCP-3^ signals (Figure 6A).

**Figure 6.**
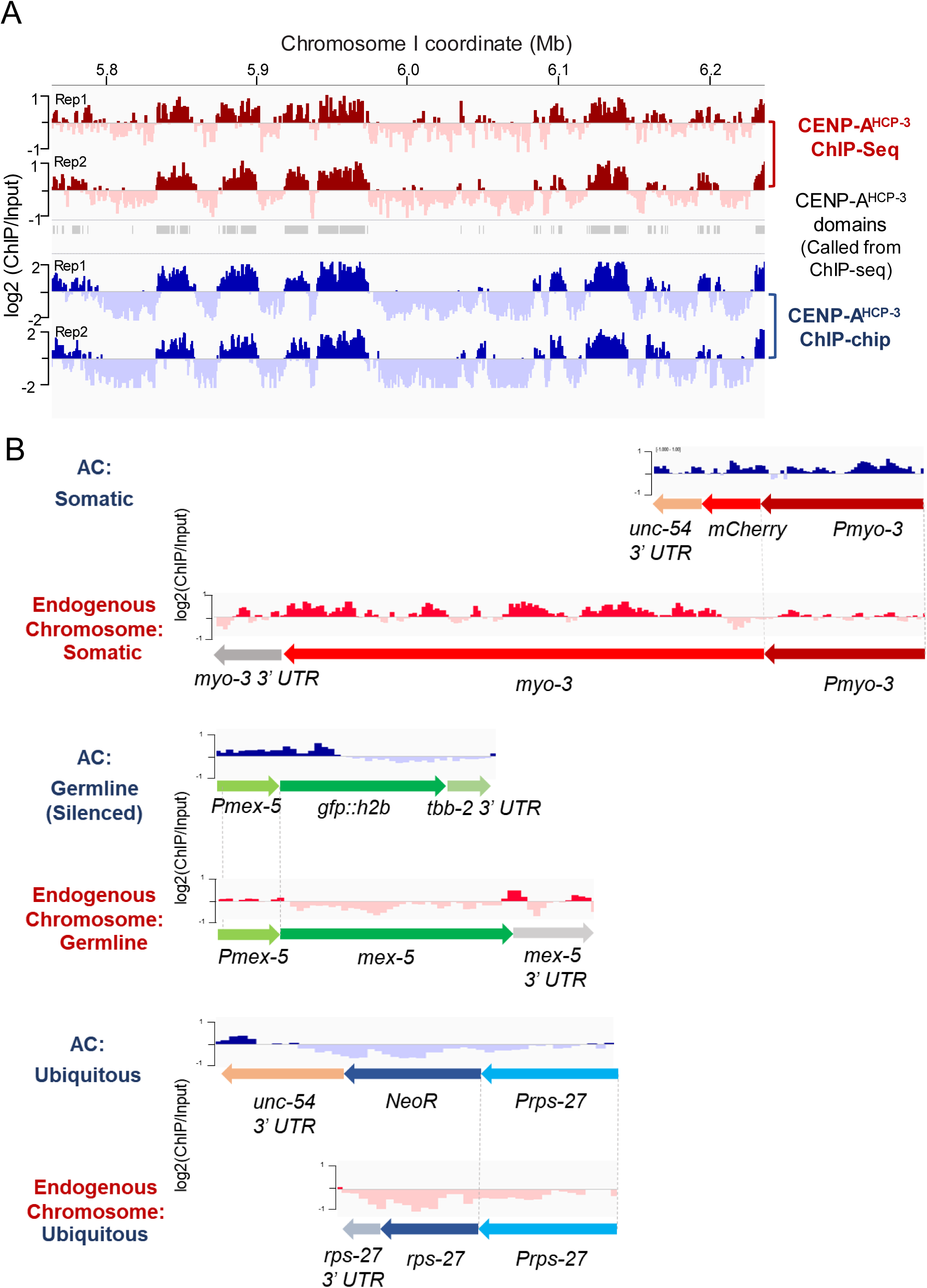
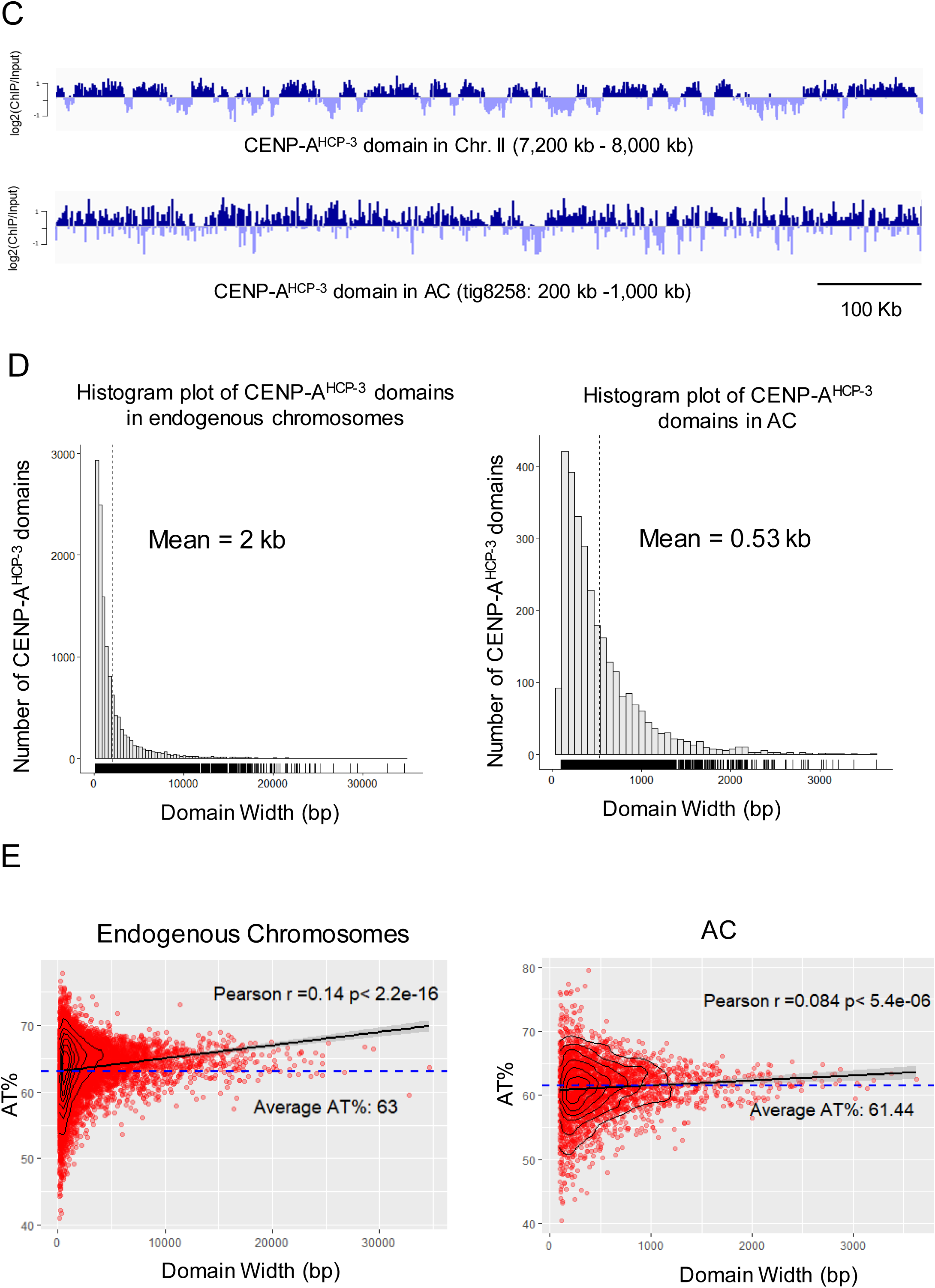
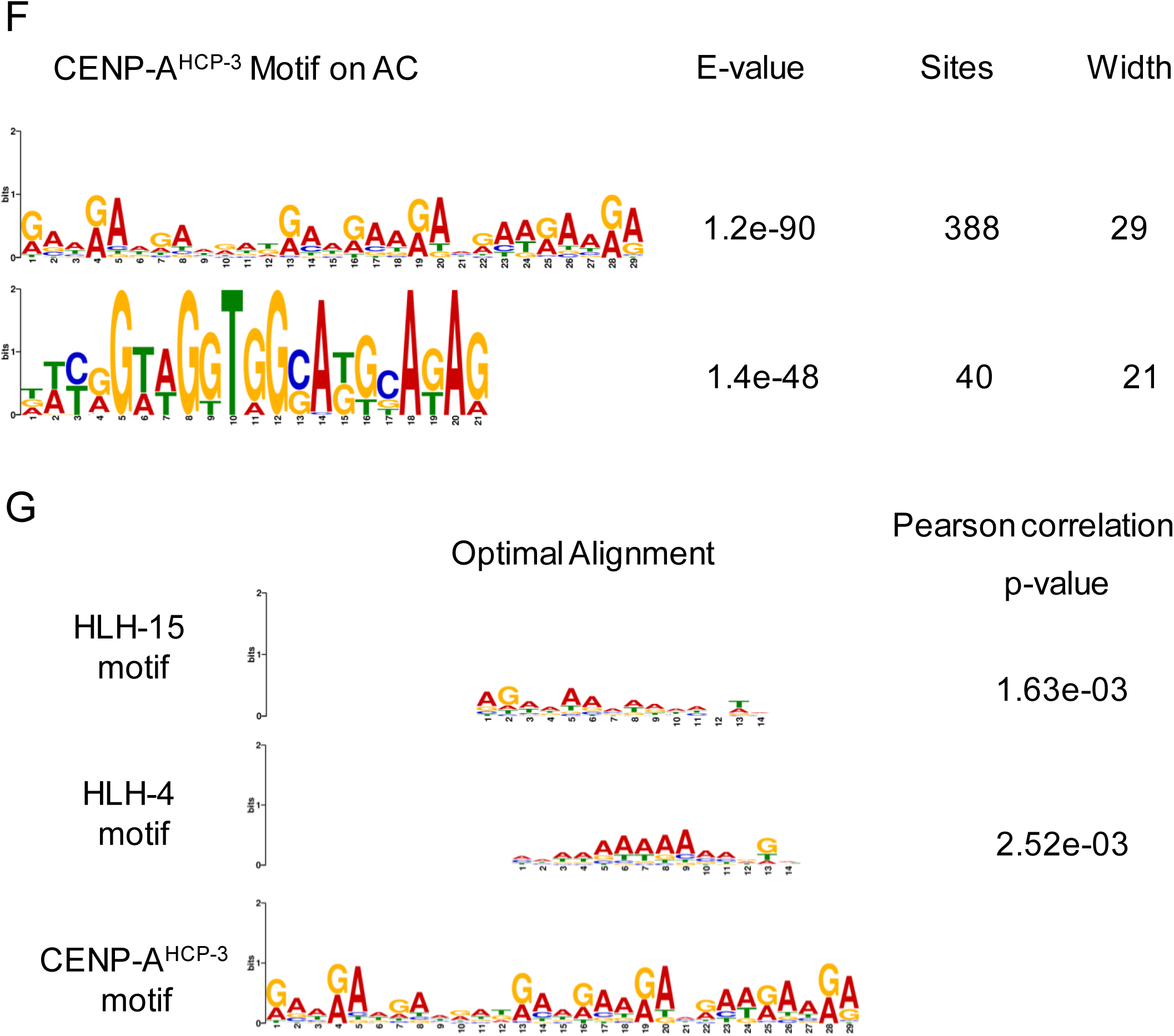
(A) Representative enrichment of CENP-A^HCP-3^ and domains on chromosome I by current ChIP-seq replicates and previous ChIP-chip replicates. (B) CENP-A^HCP-3^ localization on the AC is enriched in the somatic mCherry marker, partially excluded in silenced germline transcription regions, and excluded in ubiquitous drug marker. The corresponding endogenous genes with the same promoters were analyzed and shown in parallel. (C) Representative CENP-A^HCP-3^ domains in a 800-kb region in endogenous Chromosome II (7,200 kb - 8,000 kb) and in AC contig tig8258 (200 kb - 1,000 kb). (D) Histogram plots of CENP-A^HCP-3^ domains in endogenous chromosomes (left panel) and in AC (right panel). (E) Dot plot of AT-content (%) of each CENP-A^HCP-3^ domain against its size in endogenous chromosomes (left panel) and in AC (right panel). The linear regression line to the scatter plot model was indicated by the black line with 95% confidence region. (F) CENP-A^HCP-3^ motifs found from the centromeric domains in AC. (G) The motifs found in the UniPROBE database with a significant match to the CENP-A^HCP-3^ motif on AC.

Furthermore, we evaluated the CENP-A^HCP-3^ pattern on the complex AC. Unlike in budding yeast, in which CENP-A^CSE-4^ was only deposited on the AT-rich, 125-bp centromeric DNA, specifically on CDEII (29), CENP-A^HCP-3^ was rather uniformly deposited on the worm AC comprising of fragmented yeast sequences, suggesting that the *C. elegans* holocentromere is more likely to be epigenetically determined than sequentially determined. Besides, the CENP-A^HCP-3^-positive and CENP-A^HCP-3^-negative domains on the marker genes (average from 10-12 copies, table S4) were in consistency with the pattern on the endogenous genes driven by the same promoters. In endogenous chromosomes, CENP-A^HCP-3^ domains were generally excluded from ubiquitous, embryonic and germline expression regions (21). Consistently, in the propagated AC, CENP-A^HCP-3^ is depleted in the ubiquitous NeoR gene marker, and is enriched in the somatic mCherry marker expressed in body wall muscles, which is not expressed in germline or early embryos (Figure 6B). Because *C. elegans* germline tends to silence exogenous DNA, the germline GFP::H2B marker on this complex AC was also silenced in this strain. Interestingly, CENP-A^HCP-3^ on the germline GFP::H2B marker is half enriched (in which this half is upstream of the somatic mCherry marker, Figure 1 D) and half depleted (in which this half is downstream of the ubiquitous NeoR gene) (Figure 6B). This finding further suggests that CENP-A^HCP-3^ could be negatively regulated by the transcription level of the gene locus (Figure 6B).

### Correlation of CENP-A^HCP-3^ domain width with AT-contents and specific CENP-A^HCP-3^ motifs

We obtained a lower coverage (∼11X on average, as compared with ∼30X on endogenous chromosomes) of CENP-A^HCP-3^-enriched sequences on the AC, possibly due to the monosomic nature of the AC, or mosaicism of the AC within an animal, even under antibiotic selection. Nonetheless, we could map CENP-A^HCP-3^ domains to most of the regions on the contigs of the AC. The width of CENP-A^HCP-3^ domains on the AC is shorter and more scattered when compared with CENP-A^HCP-3^ domains on endogenous chromosomes. Representative snapshots of the CENP-A^HCP-3^ enrichment on a 800-kb region, on chromosome II (7,200 kb - 8,000 kb) and contig tig8258 (200 kb - 1,000 kb) of the AC, are shown (Figure 6C). The mean CENP-A^HCP-3^ domain width on endogenous chromosomes is 2 kb, but is 530 bp on the AC (Figure 6D). Interestingly, there is a correlation of CENP-A^HCP-3^ domain width with AT-content in the domains. However, this correlation is stronger on the endogenous chromosomes than on the AC (Figure 6E). Based on the sequences of CENP-A^HCP-3^ domains on this AC and using MEME analysis, we have identified CENP-A^HCP-3^ motifs with high A and G enrichments (Figure 6F). The most abundant motif, [GAA]_X10_, has a significant match to several transcription factor (TF) binding sites in the UniPROBE database (30), like HLH-15, HLH-4, MDL-1, HLH-2, CND-1, HLH-14, MXL-1, LIN-32, HLH-8 and HLH-19. Among them, motifs of HLH-15 (head and tail neurons expression) and HLH-4 (head, body and tail neurons expression) share the highest similarity with CENP-A^HCP-3^ motif discovered in this AC (Figure 6G).

## DISCUSSION

Repetitive and AT-rich sequences are the common features in most regional monocentromeres and some holocentromeres (31,32). In *C. elegans*, holocentromeres can rapidly establish on microinjected foreign DNA in the absence of any worm genomic DNA sequence. We used the GFP::LacI/LacO DNA tethering system to visualize newly formed artificial chromosomes. By microinjection of DNA with different AT-contents, or in repetitive or complex contexts, we tested the requirement and preference of AT-content and sequence repetitiveness in holocentromere establishment on *C. elegans* artificial chromosomes.

High AT-content sequences were routinely found in point monocentromeres, like in *S. cerevisiae*, and in many regional monocentromeres, such as in *S. pombe, Drosophila* and human (31). The *S. cerevisiae* point centromeres are strongly dependent on the high AT-content sequence (5). H3 nucleosome occupancy is known to be affected by the combination of DNA sequences and chromatin remodeling factors (33). AT-rich or poly (dA:dT) sequence may have a lower affinity for the H3 nucleosome (34), but is preferred for CENP-A nucleosome deposition (35). Studies on ectopic non-centromeric DNA regions of human neocentromeres showed that they had some similar features as in native centromeres, in which the CENP-A-binding domains on neocentromeres had slightly higher AT-contents as compared to that of the total genomic DNA (36). We hypothesized that high AT-content sequences may affect the chromatin conformation to facilitate CENP-A nucleosome deposition. In this study, we tested this hypothesis by comparing the mitotic segregation rates of newly formed ACs with different AT-contents. We found that ACs with high AT-content (74%) acquired segregation competency significantly faster than low and medium AT-content ACs (52% and 66%), suggesting that this high AT-content sequence may be more efficient in recruiting centromeric proteins than lower AT-content sequences. Genome-wide analysis of CENP-A^HCP-3^ domains from ChIP-seq also shows a correlation of CENP-A^HCP-3^ domain width with AT-rich sequences on both natural holocentromeres (endogenous) or newly established holocentromeres (on ACs), suggesting that CENP-A^HCP-3^ may tend to accumulate, or be more stable at higher AT-content regions, such that larger CENP-A^HCP-3^ domains could be formed. In this study, we demonstrated that high AT-content sequences also facilitate *de novo* centromere formation in holocentric *C. elegans*, which suggests that monocentric and holocentric centromeres may share some common DNA features and rules for centromere establishment.

Tandem repeats are the common sequence context of human centromeres (alpha satellite repeats), mice centromeres (minor satellite repeats) and fission yeast centromeres (central core repeats), and have a high competency for *de novo* centromere formation in these organisms (6,7,37,38). In the holocentric plant, *Rhynchospora*, CENP-A also localizes on centromeric repeats (32). However, in holocentric *C. elegans*, centromeric-specific repeats has not been found in endogenous chromosomes (21). To determine the effects of sequence repetitiveness on new centromere formation, AC segregation frequencies were compared between repetitive ACs and complex ACs. Yet, repetitive and complex ACs showed similar segregation rates in each embryonic stage, indicating that sequence complexity may not influence *de novo* holocentromere formation in *C. elegans*. This is consistent with the endogenous holocentromere localization in *C. elegans*, which covers 50% of the genome, mainly on non-repetitive sequences (21).

To compare the *de novo* centromere on the AC and endogenous centromeres, we generated a propagated AC in *C. elegans* made mostly out of yeast genomic DNA. We demonstrated that this AC was maintained as an independent chromosome and can align at metaphase plate and segregate with the endogenous chromosomes in mitosis (Figure 3B). However, we found that without drug selection, the average frequency of AC being inherited to the next generation is about 60%. The inheritance of this AC, as other extrachromosomal ACs, between generations does not obey Mendel’s law (9,10). We speculated that ACs may not be stably inherited in meiosis. Indeed, the Aurora B kinase, AIR-2, that localizes to the middle region of bivalent chromosomes in the diakinesis oocytes (39), is absent on the AC (Figure 3B), suggesting that the propagated AC is an univalent chromosome. Unpaired univalent chromosome tends to be missegregated, and be excluded to the polar body during female meiosis (26), which may explain why ACs are mitotically stable, but are generally lost between generations (8-10).

To investigate the DNA concatemerization events that generate this AC, we have applied the Nanopore MinION and Illumina Mi-seq platforms to sequence the whole genome (WGS) of this complex AC-containing *C. elegans* strain, followed by *de novo* assembly of the AC sequences. The *C. elegans* genome has recently been re-sequenced multiple times and polished by long-read sequencing (40,41). Tyson et al. have reported the genome of a *C. elegans* strain with integration of a multi-copy plasmid, which occupies about 2 Mbs (40). In this study, we have reported the *de novo* assembly of the genome of a *C. elegans* strain carrying an independently maintained AC, with a size of more than 10 Mbs.

Injected supercoiled DNA can form high molecular weight concatemers formed by homologous recombination, while injection of linear DNA can form HMW non-tandem ACs by random end-joining (10). Our previous study has shown that inhibition of homologous recombination (HR, by RAD-51 knockdown) or non-homologous end-joining (NHEJ, by LIG-4 knockdown) reduce AC number in embryos, indicating that both HR and NHEJ are involved in AC formation (8). We reported the sequence structure of a propagated AC formed by co-injection of a high concentration of enzyme-digested blunt-ended yeast genomic DNA and a low concentration of a selection marker and 2 differentially expressed marker genes (NGM), in a ratio of 300:1. Based on the whole genome sequencing, we suggested that non-homologous end-joining is the dominant pathway to fuse small exogenous DNA fragments (mainly < 2 kbs) to form a HMW DNA array (> 10 Mb). Double strand breaks adjacent to a region of homology can stimulate homologous recombination within 25 kb in *C. elegans* (42). On the NGM sequences, two *unc-54 3’ UTR* sequences (∼760 bp) are ∼1.6 kb apart. The homologous sequences are sufficiently long and close enough for concatemerizing through HR. However, we have not found any duplicated NGM markers that are tandemly oriented on the assembled contigs (Table S5). Instead, full-length NGM marker or its partial fragments (Table S4) were individually interspaced by yeast genomic DNA, suggesting that NHEJ dominated the fusion of microinjected foreign DNA fragments in the *C. elegans* gonad(Table S5 and Figure 5B). It is possible that NHEJ outcompetes HR when a large number of DNA fragments without HR regions are present, and only a low abundance of DNA fragments with HR regions are present. HR efficiency and usage may depend on the speed of pairing up with homologous sequences. Indeed, the inhibition of NHEJ factor increases the use of HR in CRISPR-induced double-strand break repairing in *C. elegans* (43).

We found that DNA fragments from each yeast chromosome were used in AC formation (Figure 5A and B). However, the fragmented yeast genomic DNA was not incorporated evenly in the AC forming process. Although the size of the AC is similar to that of the entire yeast genome, only a quarter of the yeast genomic sequences were chosen to be used in AC formation, which suggests that some sequences have been incorporated multiple times into the AC (Figure 5C). By aligning the restriction enzyme-digested yeast DNA fragments to the assembled AC contigs, we were able to reveal the incorporated sequences. A histogram plot of the length of the incorporated sequences shows a peak at about 1 kb (Figure 5C). Surprisingly, DNA with length below 500 bp was rarely incorporated into the AC, despite the fact that these short fragments were highly abundant in the injection mixture. In contrast, larger DNA molecules have higher chances to be incorporated in the AC. To further confirm this phenomenon, we microinjected a 300-bp linear DNA fragment with 8 copies of LacO (L8xLacO) to the *C. elegans* gonad. By live-cell time-lapse imaging, we found that microinjecting L8xLacO at the same concentration (100 ng/μl), as we had used for microinjection of L64xLacO, only generated tiny ACs (based on GFP::LacI foci sizes) that can barely segregate during mitosis, even in late embryonic stage (Figure S5B). This result suggests that microinjected DNA sequences below 500 bps was very inefficient in forming segregating ACs.

The centromeres in most organisms are located in intergenic regions. Integrating an activated gene into the centromere of *Candida albicans* (44) or tethering transcription activators to the human artificial chromosome (HAC) centromeric chromatin can disrupt centromere function (45). Previous ChIP-microarray results showed that CENP-A^HCP-3^-enriched regions in *C. elegans* embryos were excluded from the embryonic or germline transcribed regions. Ectopic expression of certain somatic genes in the *met-1* mutant germline resulted in the exclusion of CENP-A^HCP-3^ in those areas (21). These findings suggest that non-expressed regions are the preferred sites for holocentromere formation in *C. elegans*, whereas the expressed regions or previously expressed regions may contain memory markers that inhibit CENP-A^HCP-3^ deposition.

To test if this rule in endogenous holocentromeres also applies to the *de novo* holocentromere formed on ACs, we constructed plasmids that express reporter genes under different stage-specific promoters and co-injected them individually with p64xlacO for analyzing the AC segregation efficiency. However, we cannot observe the germline expression of GFP driven by an ubiquitous *his-72* promoter or a germline *pie-1* promoter (data not shown). Ectopic genes, especially those on repetitive ACs, were commonly suppressed in *C. elegans* germline by chromatin remodeling factors and RNAi factors (46). We instead examined the CENP-A^HCP-3^ localization on a complex, propagated AC by ChIP-seq. Our ChIP-seq results on *C. elegans* genome shows a high correlation between replicates and with previous ChIP-microarray results (21) (Figure 6A, S6A and S6B). The antibiotics resistance gene *NeoR*, driven by ubiquitous *rps-27* promoter, shows a negative CENP-A^HCP-3^ signal. The endogenous *rps-27* gene also shows a negative CENP-A^HCP-3^ pattern. Similarly, the mCherry gene driven by somatic *myo-3* promoter shows a positive CENP-A^HCP-3^ signal, and this is also consistent with the enrichment of CENP-A^HCP-3^ found on the endogenous *myo-3* promoter and its downstream coding gene body (Figure 6B). The *GFP::H2B*, driven by germline *mex-5* promoter on this propagated AC, were silenced and not detectable under fluorescent microscopy. Interestingly, CENP-A^HCP-3^ signal is positive on the promoter and the first half of the GFP coding region (upstream of the somatic marker), but is negative on the other half of the GFP and the H2B coding region (close to the downstream ubiquitous NeoR gene). These results suggest that CENP-A^HCP-3^ could spread to the silenced germline genes, but the CENP-A^HCP-3^ expansion may be inhibited by the nearby transcriptionally active region (Figure 6B). Besides, the CENP-A^HCP-3^ distribution on 3’ UTRs are more dependent on the activity of their upstream genes, on both the endogenous chromosome and the AC (Figure S6C). Therefore, we conclude that CENP-A^HCP-3^ on *de novo* holocentromere can also be excluded by ubiquitous transcription, and the unexpressed regions are preferred for *de novo* holocentromere formation.

The fragmented yeast sequences do not contain many full genes, and whether any of the yeast genes are expressed in worms is currently unknown. However, the correlation of CENP-A^HCP-3^ domain sizes in both endogenous chromosomes and the AC suggest that AT-rich sequences are preferred (Figure 6D and E), and this is consistent with other regional centromeres and holocentromeres (31,32). The most abundant motif we found from the CENP-A^HCP-3^-enriched AC domains is a 29-bp [GAA]_X10_ sequence (Figure 6F). Interestingly, by comparing with the protein motif database, several HLH transcription factors, like HLH-15 and HLH-4, have high correlation with the [GAA]_X10_ motif (Figure 6G), which is similar to the enrichment of transcription factor hotspots (HOT sites), like HLH-1 motif, found in CENP-A^HCP-3^ single-nucleosome peaks on endogenous chromosomes (47,48). These data suggest that sequences preferred for HLH TFs binding is potentially facilitating *de novo* CENP-A^HCP-3^ nucleosome deposition. Furthermore, the [GAA]_X10_ motif also has a potential to form non-B-form DNA conformation (49), consistent with non-B-form DNA commonly found in mammalian centromeres and its high competency of *de novo* centromere formation in human cells (50,51). Altogether, these findings indicate that the AT-sequence, non-repetitive and non-transcriptional status preference, and [GAA]_X10_ motif enrichment of de novo centromere formation at ACs are consistent with holocentromere features of endogenous *C. elegans* chromosomes. These results can validate the use of *de novo* centromere formation on ACs to simulate the events that occur in centromere formation of endogenous chromosomes.

Overall, this holocentric model organism system, amenable to foreign DNA injection and AC formation with *de novo* holocentromere formation, allows us to manipulate the input DNA, compare the *de novo* centromere formation efficiency by live-cell imaging, and analyze the positions of CENP-A^HCP-3^ domains in the propagated AC. We have compared different epigenetic and DNA sequence parameters in order to elucidate the preferences and their respective functions in *de novo* holocentromere formation in this *in vivo*, real-time system.

## ACKNOWLEDGEMENT

We thank J Hui, W Nong, J Dumont, KM Chan, R Ng, Y Zhai, and W Den for critical reading of earlier versions of the manuscript, and the Yuen lab for discussion.

