## Supplementary_table&figures_legends for "DNA Sequence Preference for *De Novo* Centromere Formation on a *Caenorhabditis elegans* Artificial Chromosome"

**SUPPLEMENTARY TABLE LEGENDS AND FIGURE LEGENDS**

Table S1. Summary of MinION sequencing data from two flow-cells.

Table S2. Comparison of mappable MinION reads to the worm reference genome (WS245) based on All reads and Pass reads.

Table S3. Summary of genome assemblies and polishing from a *C. elegans* strain (WYY35) carrying an artificial chromosome, based on different genome assembly pipelines using MinION, Mi-seq and combined datasets.

Table S4. The copy number of marker genes in the AC.

Table S5. The localization of marker genes in the AC.

Figure S1. Whole-genome sequencing of a *C. elegans* strain with a complex artificial chromosome L2.1 (WYY35). Raw data produced from MinION flow-cells were base-called by Metrichor. (A, B) Histogram of the read numbers was plotted against the read length yielded from each flow-cell. The blue line indicates the N50 of read length. (C, D) Histograms of read length and Phred quality were shown. The number of reads from each flow-cell in different read lengths and read qualities were indicated by colors.

Figure S2. MinION individual read accuracies and read coverage when aligned to the reference worm genome (WS245). (A) Graphmap aligner was used for mapping. Read base-called qualities were plotted against mapped read percent identity. Most of the reads have a percent identity of about 85-90%, but with a long tail to identities of ~60%. Pearson correlation: r = 0.75, p-value = 0. A dashed red circle shows a small portion of high-quality reads with low mapping identity, indicating these reads might not belong to the worm genomic DNA. (B) Alignment percentage read identity was plotted against aligned read length. This is a comparison of mapping quality (based on percent identity) for different read lengths after alignment using Graphmap aligner. No length bias was observed, with long reads having the same percentage identity as short ones. Pearson correlation: r = 0.04, p-value = 0. (C) Coverage for *C. elegans* reference genome (WS245) based on all reads generated from 2 MinION flow-cells. (D) Coverage for *C. elegan*s reference genome (WS245) based on Pass reads, filtered base on Phred quality (>q6).

Figure S3. Mummer alignments of assembled contigs from MinION reads to *C. elegans* reference genome (WS245). The dot plot displays the one-to-one mapping between the assembled contigs sets (on the Y-axis) and the reference chromosomes (on X-axis). Forward matches are indicated in red and reverse complement matches in blue.

Figure S4. Misassembled NGM and yeast sequences to a worm contig by Canu direct assembly.

(A) A schematic of yeast sequence fragments and the co-injection markers (*Pmyo-3::mCherry:: unc-54 3' UTR* and *Pmex5::GFP::tbb-2 3' UTR*) on tig93. DNA fragments belong to individual yeast chromosomes and NGM marker genes are indicated by different colors. (B) The misassembled region has low read coverage, which was indicated by a red arrow. MinION reads were re-mapped to the AC contigs from direct Canu assembly.

Figure S5. (A) Self-alignment of the largest contig, tig8258, of the assembled AC shows that some short sequences are incorporated multiple times. The co-injection markers NGM (*Prps-27::NeoR::unc-54 3' UTR*; *Pmex5::GFP::tbb-2 3' UTR*; *Pmyo-3::mCherry:: unc-54 3' UTR*) were interspaced in between yeast genomic DNA on tig8258. DNA fragments aligned to individual yeast chromosomes and NGM markers are indicated by different colors. (B) The tiny ACs formed by microinjection of 8xLacO has no segregation competency. Linear 8xLacO (315 bp) fragments were microinjected at a concentration of 100 ng/μl. No AC segregation was observed even in the late embryo stage.

Figure S6. Comparison of CENP-A$^{HCP-3}$ on endogenous chromosomes by ChIP-seq in this study and ChIP-chip from a previous study (21). Pairwise scatter plotted by (A) Pearson correlation method and (B) heatmap plotted by Spearman correlation method both show that the localization of CENP-A$^{HCP-3}$ domains on endogenous chromosomes detected by ChIP-seq is comparable to previous ChIP-chip results. The correlation coefficient in Pearson correlation and Spearman correlation between ChIP-seq replicates are 0.83 and 0.84, respectively. The cross-platform correlation between pairs of replicates of ChIP-chip and ChIP-seq are higher than 0.81. (C) Comparison of CENP-A$^{HCP-3}$ distribution on the same 3' UTRs in the AC and endogenous chromosomes.
