## Supplementary_table&figures for "DNA Sequence Preference for *De Novo* Centromere Formation on a *Caenorhabditis elegans* Artificial Chromosome"

Table S1. Comparison of the assembly of the AC by directly assembling the whole genome (endogenous chromosomes and the AC), or by assembling reads that are filtered out (excluded) from worm genomic sequences, as outlined in Figure 4.

| Flow-cell ID | FAF03915 | | FAF03410 | |
|---|---|---|---|---|
| | Total | Filtered | Total | Filtered |
| **Read Count** | 636360 | 483360 | 805812 | 640037 |
| **Total Yield** | 4.92 G bases | 3.92 G bases | 5.03 G bases | 4.18 G bases |
| **Sequence Length - Average** | 7.74 K bases | 8.10 K bases | 6.24 K bases | 6.53 K bases |
| **Sequence Length - Median** | 7.48 K bases | 7.48 K bases | 5.78 K bases | 5.78 K bases |
| **Sequence Length - Mode** | 6.82 K bases | 7.70 K bases | 578.00 bases | 578.00 bases |
| **Longest Read** | 131.78 K bases | 93.50 K bases | 149.69 K bases | 121.02 K bases |
| **QScore – Average** | 7.1 | 7.8 | 7.2 | 7.8 |
| **QScore – Median** | 7.5 | 7.9 | 7.6 | 7.9 |
| **QScore – Mode** | 8.4 | 8.4 | 8.4 | 8.4 |

Table S2. Comparison of mappable MinION reads to the worm reference genome (WS245) based on from All reads and Pass reads.

| | **All read [a] coverage and identity** | **Pass read [b] coverage and identity** |
|---|---|---|
| **Reference genome size (WS245)** | 100,286,401 | |
| **Number of reads** | 1,442,348 | 1,081,581 |
| **Mapped reads** | 1,142,250 / 79.19% | 957,042 / 88.49% |
| **Unmapped reads** | 300,098 / 20.81% | 124,539 / 11.51% |
| **Read min/max/mean length** | 5 / 149,685 / 6,902.73 | 5 / 95,097 / 7,363.16 |
| **Duplicated reads (estimated)** | 32,252 / 2.24% | 23,280 / 2.15% |
| **Duplication rate** | 1.70% | 1.52% |
| **Clipped reads** | 875,288 / 60.68% | 741,107 / 68.52% |
| **Mean Mapping Quality** | 34.81 | 35.13 |
| **Mean Coverage** | 88.8253 | 75.8705 |
| **Coverage Standard Deviation** | 135.5784 | 109.2802 |
| **General error rate** | 20.73% | 18.87% |
| **Mismatches** | 1,568,580,609 | 1,218,756,648 |
| **Insertions** | 192,568,865 | 147,374,109 |
| **Mapped reads with at least one insertion** | 99.99% | 100% |
| **Deletions** | 694,098,290 | 553,275,323 |
| **Mapped reads with at least one deletion** | 100% | 100% |
| **Homopolymer indels** | 37.76% | 39.32% |

[a] : All reads produced from two MinION nanopore flow-cells.
[b] : Pass reads are high-quality reads isolated when the Phred quality is > 6.

Table S3. Summary of genome assemblies and polishing from a *C. elegans* strain carrying an artificial chromosome, based on different genome assembly pipelines using MinION, Mi-seq and combined datasets.

| | **MinION Canu (All reads[a])** | **MinION Canu (Pass reads[b])** | **Mi-seq SPAdes** | **Canu (All) + Pilon (x3) [c]** |
|---|---|---|---|---|
| **Total Bases** | 9.95 G bases | 8.1G bases | 5.4 G bases | 9.95 G bases + 5.4 G bases |
| **Contigs** | 241 | 252 | 38,187 | 241 |
| **Total Assembly (bp)** | 114,955,672 bp | 115,248,067 bp | 110,413,755 bp | 119,160,402 bp |
| **N50** | 1,487,696 bp | 1,404,122 bp | 22,768 bp | 1,515,988 bp |
| **Longest contig** | 4,707,476 bp | 4,862,513 bp | 188,252 bp | 4,774,338 bp |
| **Identity to reference** | 95.84% | 95.84% | 99.94% | 99.79% |
| **Aligned bases (REF vs. QRY )** | 100139477 (99.85%) vs. 99505683 (86.46%) | 100173646 (99.89%) vs. 99624555 (86.44%) | 100183478 (99.90%) vs. 98876593 (91.17%) | 100147804 (99.86%) vs. 103389851 (86.77%) |

[a]: All reads produced from two MinION nanopore flow-cells.
[b]: Pass reads are high-quality reads isolated when the Phred quality > 6.
[c]: Assembled contigs using All reads by Canu were polished three rounds by Pilon.

Table S4. The copy number of marker genes in the AC.

| Type | Query name | Length | Hits | Query cover |
|---|---|---|---|---|
| full length gene | Pmex5::GFP::tbb-2 3'UTR | 2,136 | 10 | 100.00% |
| full length gene | Pmyo-3::mCherry::unc-54 3'UTR | 4,197 | 12 | 99.90% |
| full length gene | Prps-27::NeoR::unc-54 3'UTR | 2,288 | 11 | 99.96% |
| gene cassette | Pmyo-3 | 2,503 | 12 | 100.00% |
| gene cassette | Pmex-5 | 486 | 13 | 100.00% |
| gene cassette | Prps-27 | 797 | 15 | 99.87% |
| gene cassette | GFP | 867 | 11 | 100.00% |
| gene cassette | mCherry | 864 | 19 | 100.00% |
| gene cassette | NeoR | 785 | 13 | 100.00% |
| gene cassette | Unc-54 3'UTR | 760 | 29 | 100.00% |
| gene cassette | Tbb-2 3'UTR | 332 | 10 | 100.00% |

Figure S1

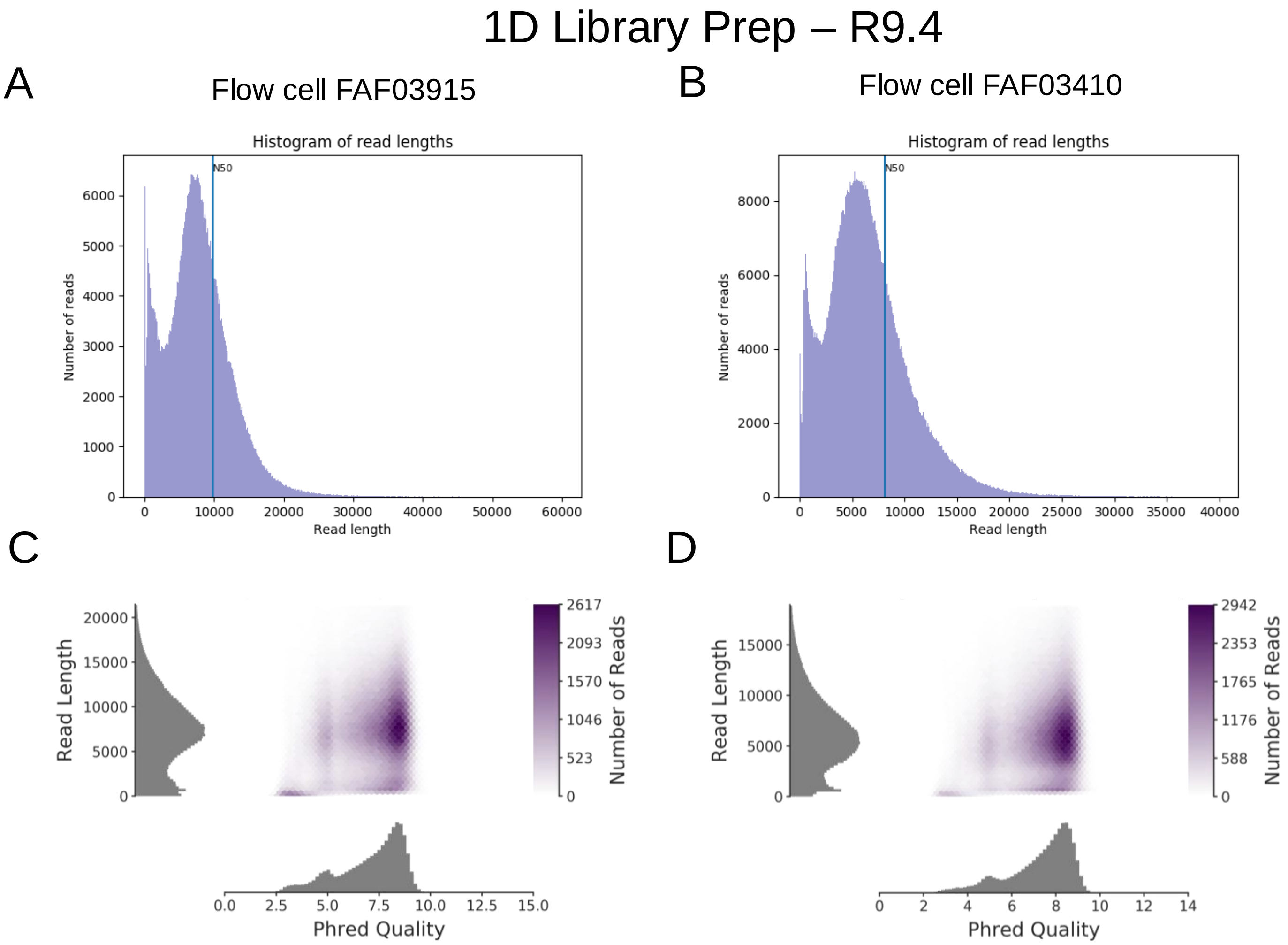
1D Library Prep – R9.4
A
Flow cell FAF03915
Histogram of read lengths
N50
Number of reads
Read length
B
Flow cell FAF03410
Histogram of read lengths
N50
Number of reads
Read length
C
Read Length
Number of Reads
Phred Quality
D
Read Length
Number of Reads
Phred Quality

Figure S2

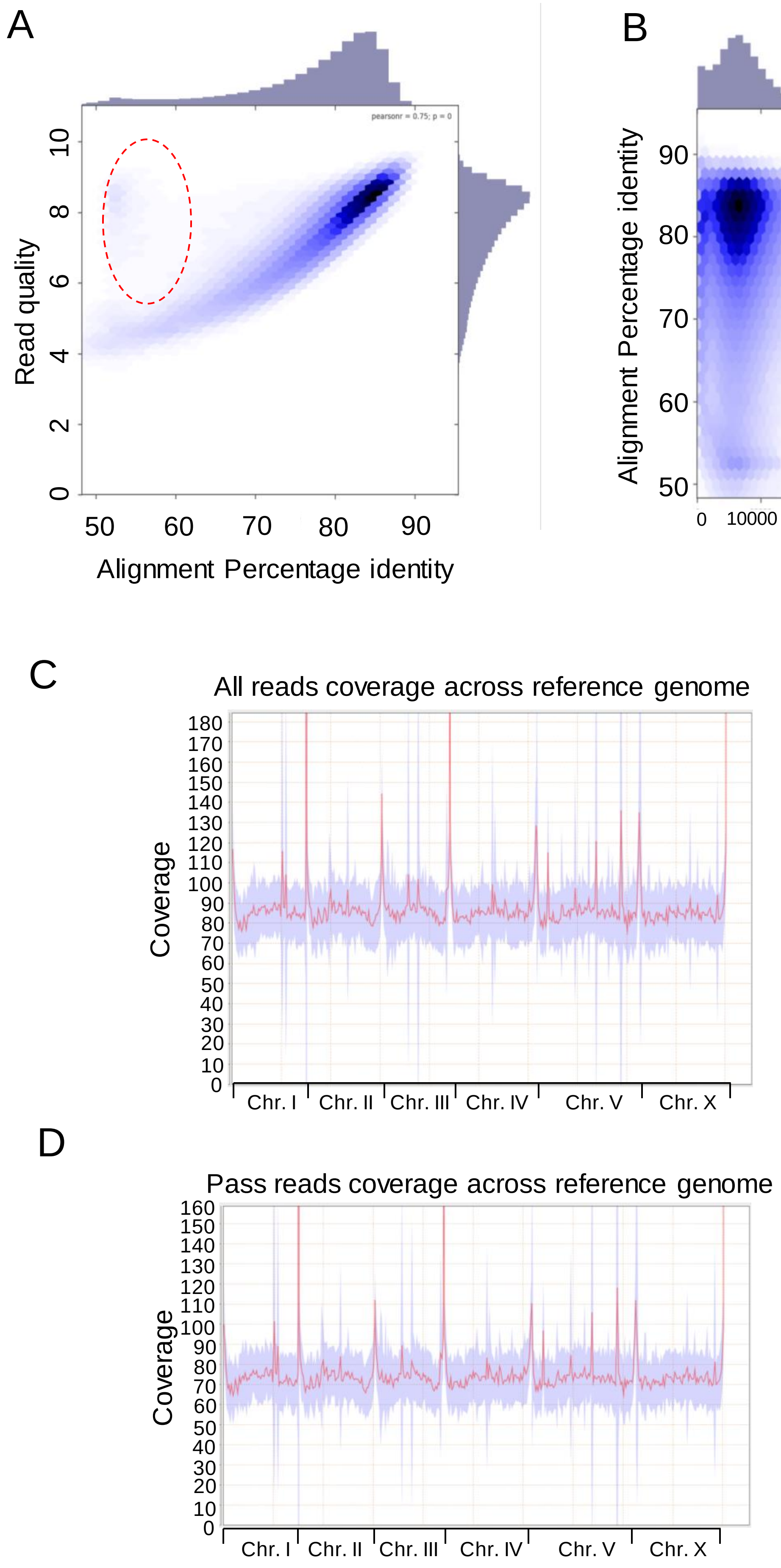

A
pearsonr = 0.75; p = 0
Read quality
0
2
4
6
8
10
50
60
70
80
90
Alignment Percentage identity
B
pearsonr = 0.04; p = 0
Alignment Percentage identity
50
60
70
80
90
0
10000
20000
30000
40000
50000
Aligned read length
C
All reads coverage across reference genome
Coverage
180
170
160
150
140
130
120
110
100
90
80
70
60
50
40
30
20
10
0
Chr. I
Chr. II
Chr. III
Chr. IV
Chr. V
Chr. X
D
Pass reads coverage across reference genome
Coverage
160
150
140
130
120
110
100
90
80
70
60
50
40
30
20
10
0
Chr. I
Chr. II
Chr. III
Chr. IV
Chr. V
Chr. X

### Figure S3

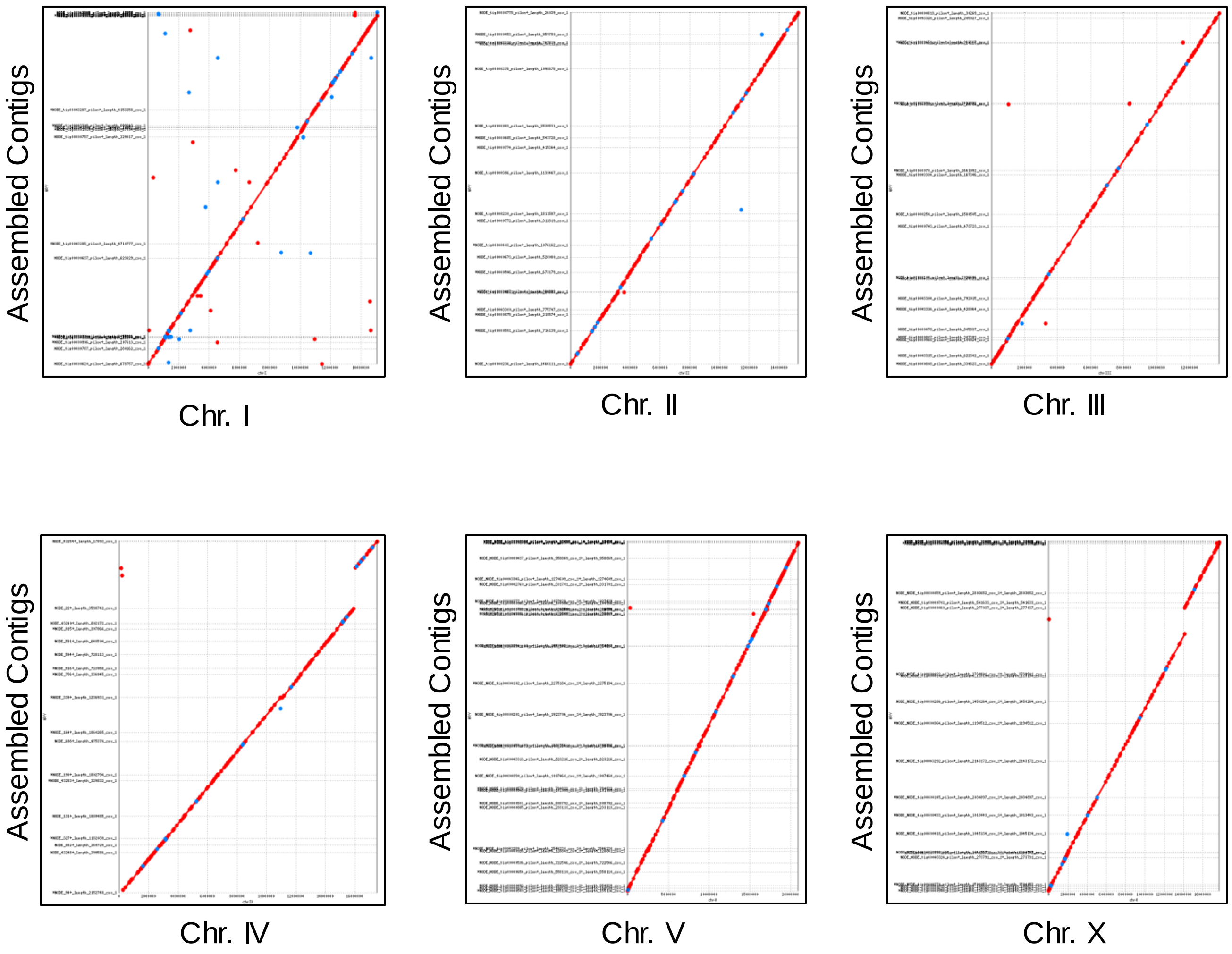

Assembled Contigs
Chr. I
Assembled Contigs
Chr. Ⅱ
Assembled Contigs
Chr. Ⅲ
Assembled Contigs
Chr. Ⅳ
Assembled Contigs
Chr. V
Assembled Contigs
Chr. X

### Figure S4

A

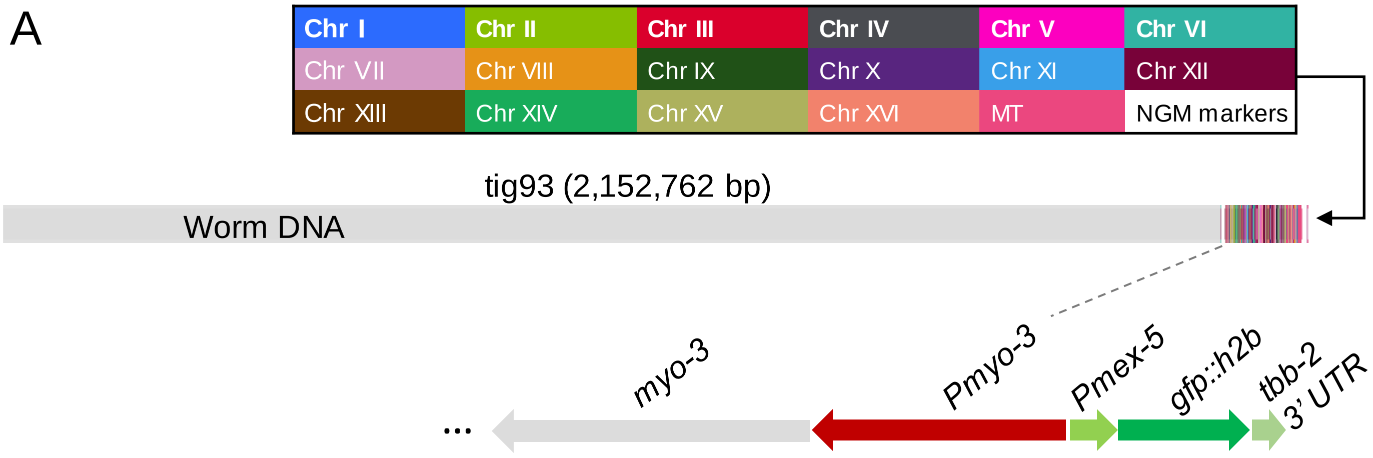
Chr I
Chr II
Chr III
Chr IV
Chr V
Chr VI
Chr VII
Chr VIII
Chr IX
Chr X
Chr XI
Chr XII
Chr XIII
Chr XIV
Chr XV
Chr XVI
MT
NGM markers
tig93 (2,152,762 bp)
Worm DNA
myo-3
Pmyo-3
Pmex-5
gfp::h2b
tbb-2
3' UTR

B

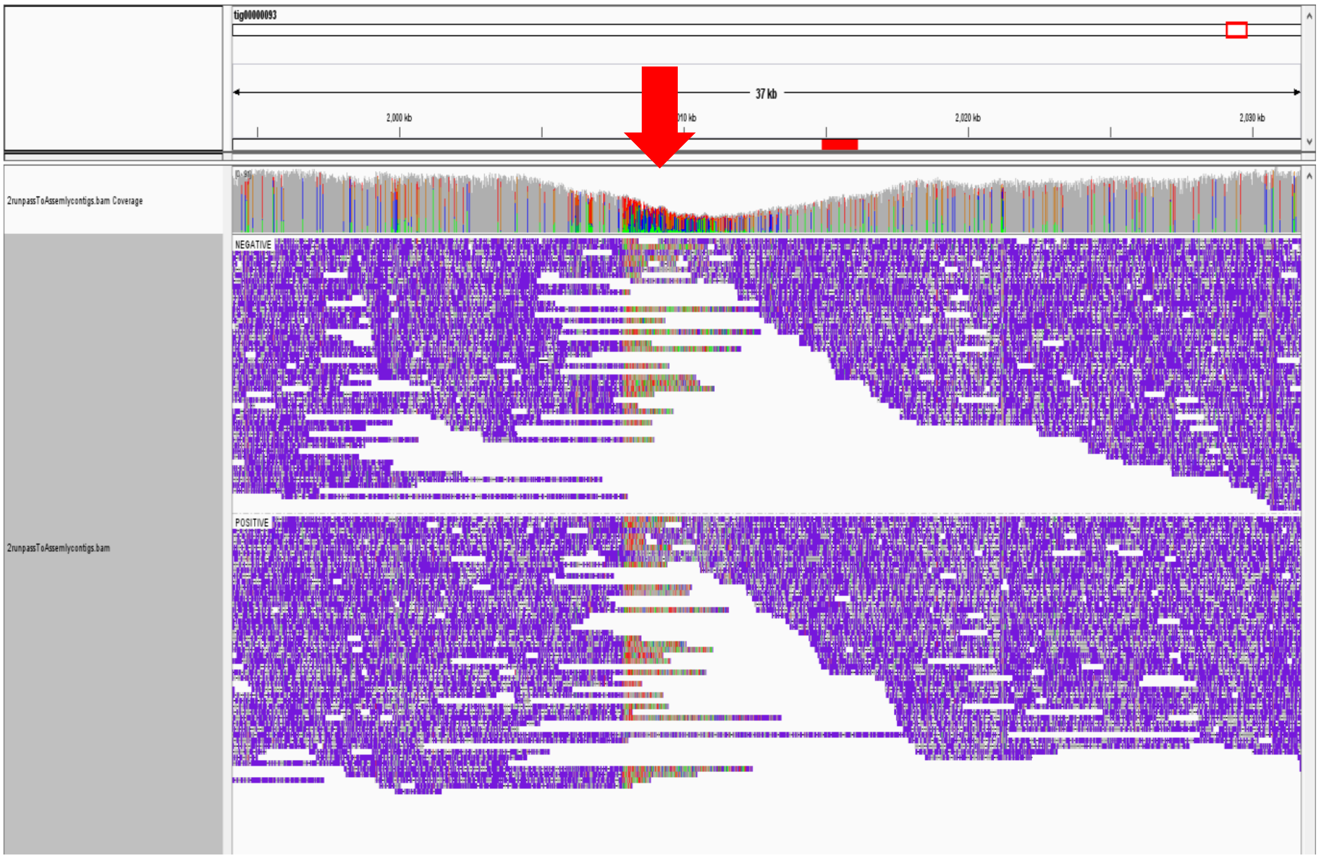

Figure S5

A

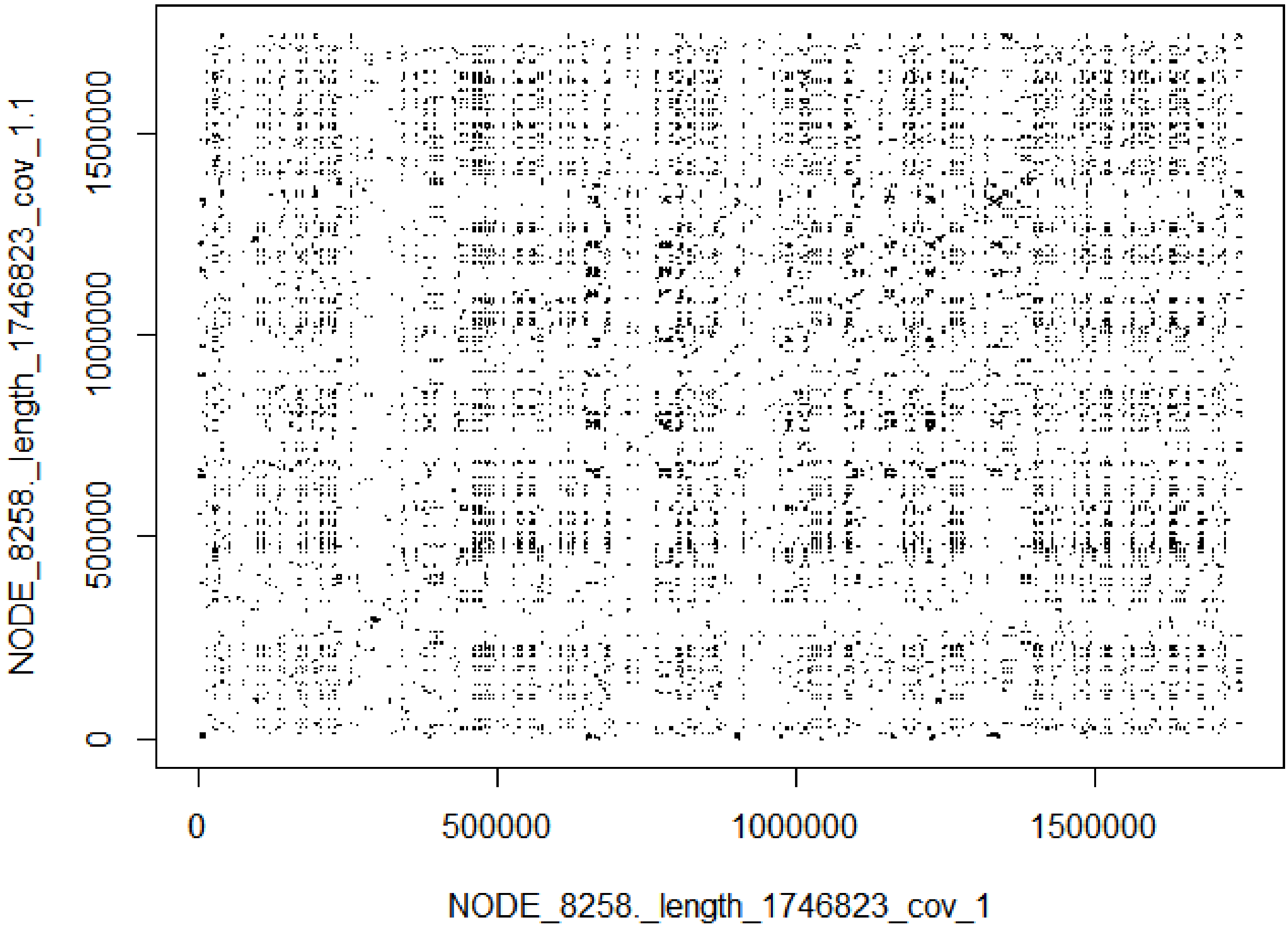
NODE_8258._length_1746823_cov_1.1
0
500000
1000000
1500000
0
500000
1000000
1500000
NODE_8258._length_1746823_cov_1

B

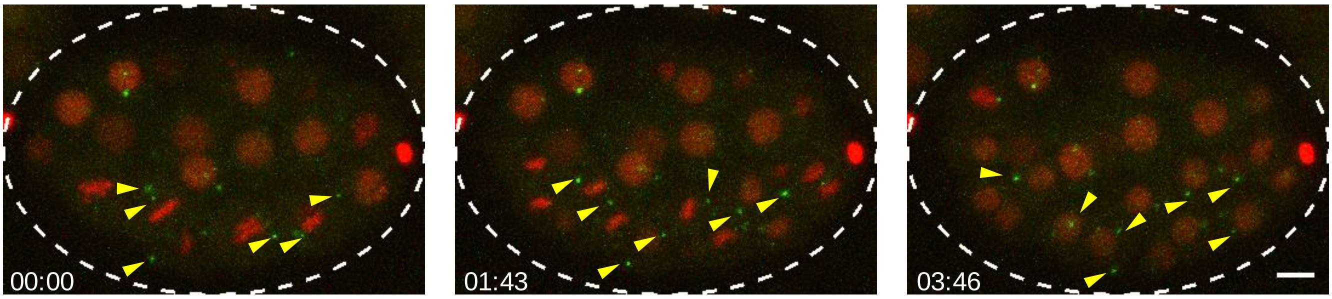
00:00
01:43
03:46

### Figure S6

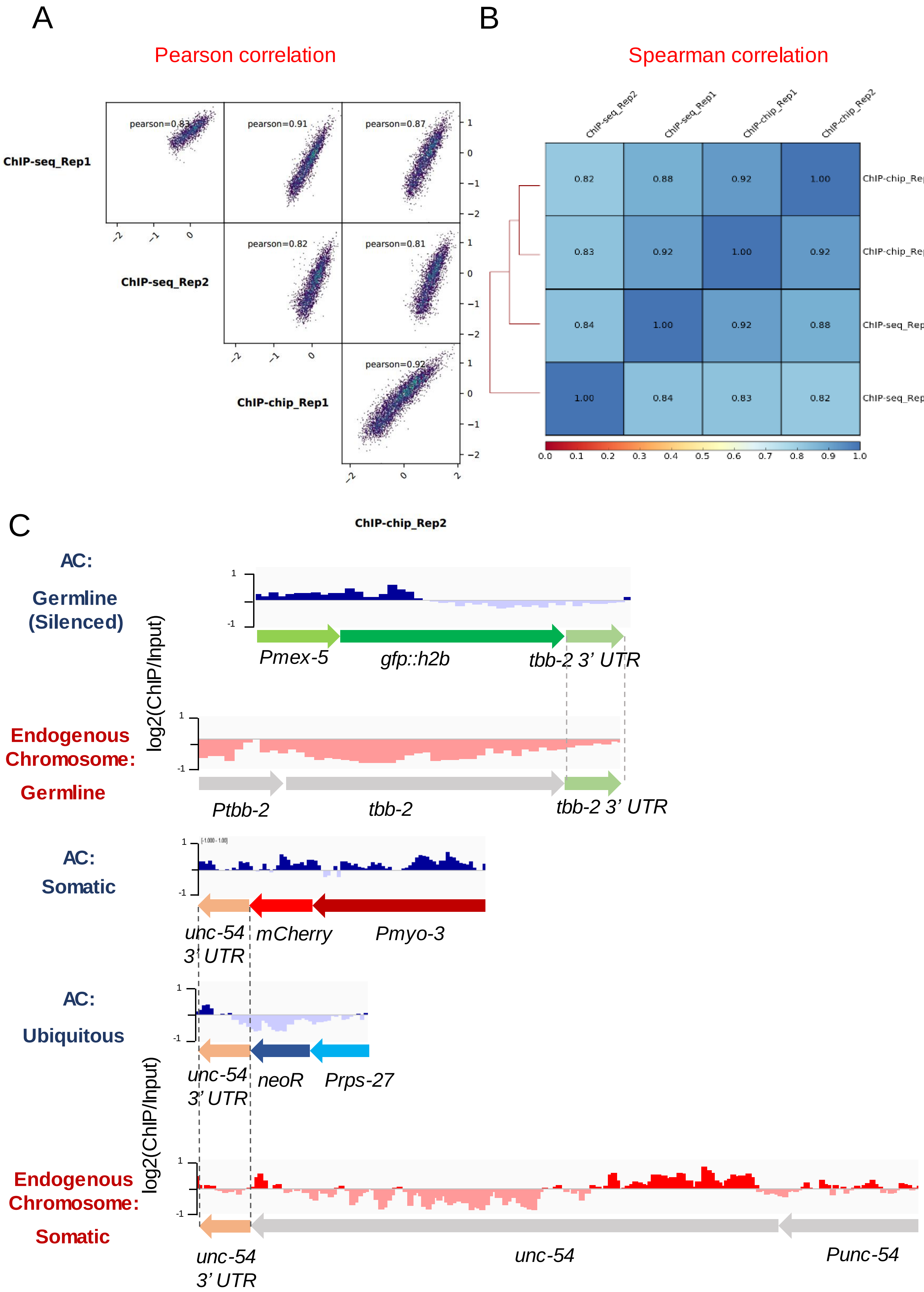

A
B
Pearson correlation
Spearman correlation
ChIP-seq_Rep1
ChIP-seq_Rep2
ChIP-chip_Rep1
ChIP-chip_Rep2
pearson=0.83
pearson=0.91
pearson=0.87
pearson=0.82
pearson=0.81
pearson=0.92
ChIP-seq_Rep2
ChIP-seq_Rep1
ChIP-chip_Rep1
ChIP-chip_Rep2
0.82 0.88 0.92 1.00
0.83 0.92 1.00 0.92
0.84 1.00 0.92 0.88
1.00 0.84 0.83 0.82
0.0 0.1 0.2 0.3 0.4 0.5 0.6 0.7 0.8 0.9 1.0
C
AC:
Germline (Silenced)
log2(ChIP/Input)
Pmex-5
gfp::h2b
tbb-2 3' UTR
Endogenous Chromosome:
Germline
Ptbb-2
tbb-2
tbb-2 3' UTR
AC:
Somatic
unc-54 3' UTR
mCherry
Pmyo-3
AC:
Ubiquitous
unc-54 3' UTR
neoR
Prps-27
Endogenous Chromosome:
Somatic
log2(ChIP/Input)
unc-54 3' UTR
unc-54
Punc-54
